# CTR1-mediated copper uptake orchestrates metabolic-epigenetic regulation of pathogenic T_H_17 cells in autoimmune disease

**DOI:** 10.64898/2026.09.22.747582

**Authors:** Lucile Noyer, Liwei Wang, Miki Jishage, Maxwell McDermott, Aidan T Pezacki, Anthony Y Tao, Ravichandran Ramasamy, Matthias Stadtfeld, Christopher J Chang, Stefan Feske

## Abstract

Pathogenic T helper 17 (pT_H_17) cells are a subset of CD4^+^ T cells driving autoimmune diseases including multiple sclerosis (MS). Compared to homeostatic T_H_17 cells and other T_H_ subsets, pT_H_17 have enhanced mitochondrial function and oxidative phosphorylation (OXPHOS) that supports their differentiation and pathogenic function. Here we identify Copper Transporter 1 (CTR1), encoded by *Slc31a1,* as essential for copper uptake in CD4^+^ T cells, OXPHOS and pT_H_17 cell differentiation and function. While copper levels are known to be higher in the cerebrospinal fluid of patients with MS compared to healthy individuals, and excess copper contributes to oligodendrocyte loss in murine models of MS, the effect of copper on T cell function and pathogenicity in MS are unclear. We demonstrate that deletion of *Slc31a1* in CD4^+^ T cells decreased intracellular copper levels, disrupting mitochondrial respiration and rewiring metabolism. These changes altered the epigenetic landscape of pT_H_17 cells by impairing DNA demethylation capacity, leading to hypermethylated DNA and altered chromatin accessibility at key binding sites for AP-1 transcription factors essential for pT_H_17 differentiation. As a result, CTR1-deficient T cells showed defective differentiation into pT_H_17 cells, with decreased production of IL-17A and expression of T_H_17 signature genes, while the differentiation of other CD4^+^ T cell subsets remained largely unaffected. Moreover, T cell-specific deletion of *Slc31a1* protected mice from central nervous system (CNS) inflammation in the experimental autoimmune encephalomyelitis (EAE) model of MS by suppressing clonal expansion of autoreactive CD4^+^ T cells. These findings establish copper as a critical regulator of pT_H_17 differentiation and function, revealing a previously unknown molecular link between copper homeostasis, metabolism and epigenetic regulation governing pT_H_17-mediated autoimmunity.

## INTRODUCTION

CD4^+^ T helper (T_H_) cells are central players in adaptive immune responses, comprising of different subsets with distinct cytokine expression and effector functions. Among these subsets, T_H_17 cells were initially defined by their production of the pro-inflammatory IL-17A/F cytokines but were later recognized to be a heterogenous subset with diverse physiological and pathological roles^1^. While homeostatic T_H_17 (homT_H_17) cells contribute to maintaining tissue homeostasis at barrier sites, pathogenic T_H_17 (pT_H_17) cells are involved in the pathophysiology of many inflammatory and autoimmune diseases^2^. Considerable effort has been directed toward characterizing pT_H_17 cells to inform the development of novel therapeutic strategies for the treatment of autoimmune diseases such as multiple sclerosis (MS), rheumatoid arthritis (RA), inflammatory bowel disease and psoriasis^3^.

Ion channels and transporters (ICTs) are transmembrane proteins that mediate the movement of ions and a broad range of metabolites across biological membranes. Several ICTs were shown to play essential roles in T cell activation, differentiation and/or function^3,4^. Among them, the calcium channel ORAI1 and the potassium channel Kv1.3 promote neuroinflammation in EAE^5,6^. To identify novel ICTs that promote autoimmune CNS inflammation and pT_H_17 cell function, we developed a targeted forward genetic screen in CD4^+^ T cells using the EAE mouse model of MS. We identified several new regulators of pT_H_17 cells in EAE including the zinc transporter ZIP3, encoded by *Slc39a3,* and the Chloride Nucleotide-Sensitive Channel 1A (CLNS1A)^7,8^. Here we report the identification of Copper Transporter 1 (CTR1), encoded by *Slc31a1*, as a critical regulator of pT_H_17 function and autoimmunity. CTR1 is a high-affinity copper importer located in the plasma membrane^9–14^. Copper is an essential micronutrient involved in key biological processes by serving as a cofactor for enzymes such as Superoxide Dismutase 1 (SOD1) and Cytochrome C Oxidase (COX)^15^. The COX complex, or complex IV, is the terminal enzyme complex of the electron transport chain (ETC). It contains two copper sites, Cu_A_ and Cu_B_, binding a total of three copper ions, essential for electron transfer during respiration^16^. Several studies reported increased copper levels in autoimmune diseases such as MS^17^, RA^18^, psoriasis^19^ and lupus^20^, suggesting a role for copper in autoimmune inflammation. Further, treatment with the copper chelator tetrathiomolybdate (TTM) protected mice from CNS autoimmunity and inflammation in the CD4^+^ T cell dependent EAE model of MS^21^, suggesting a role for copper in the pathogenicity of encephalitogenic T cells. While copper was shown to be essential to sustain the inflammatory response of macrophages^22^, and the maintenance of regulatory T cells (T_reg_)^23^, the role of copper in T_H_ cells, in particular pT_H_17 cells and autoimmunity, has not been elucidated.

Here we demonstrate that copper and CTR1 control the differentiation of pT_H_17 cells, while having lesser effects on other T_H_ subsets, by regulating their metabolism and establishing the epigenetic marks that control the expression of the pT_H_17 program. T cell specific deletion of *Slc31a1* decreases intracellular copper levels, impairs mitochondrial respiration and rewires T cell metabolism. The ensuing metabolic imbalance is associated with the accumulation of metabolites known to be potent inhibitors of TET (Ten-eleven translocation) enzymes and histone demethylases (HDM) responsible for the demethylation of DNA and histones, respectively. Together with the depletion of α-ketoglutarate, the substrate for the demethylation reaction, CTR1 deficiency resulted in DNA hypermethylation in pT_H_17 cells. The removal of DNA methylation is critical for the recruitment of AP-1 transcription factors and enforcement of the pT_H_17 transcriptional program. As a consequence, deletion of CTR1 in T cells strongly protects mice from EAE. Here, we propose a model wherein CTR1 and copper regulate pT_H_17 function through mitochondrial respiration and epigenetic regulation of key transcriptional programs. Our findings provide evidence for a copper-dependent link between metabolism and epigenetic regulation of pT_H_17 cell function that is essential for autoimmune CNS inflammation.

## RESULTS

### The copper transporter CTR1 (SLC31A1) is essential for pT_H_17 cells in EAE

To identify novel ion channels and transporters (ICT) regulating encephalitogenic pT_H_17 cells, we performed an ICT-targeted forward genetic screen using the EAE mouse model of MS as previously described^7,8^. We identified CTR1 (encoded by the murine *Slc31a1* gene) among the top 44 novel ICTs depleted in encephalitogenic pT_H_17 cells in the CNS of mice (**Fig. 1A**). To confirm the importance of CTR1 in pT_H_17 cells in EAE, we transduced 2D2 CD4^+^ T cells expressing a myelin oligodendrocyte glycoprotein (MOG)-specific T cell receptor (TCR) with either shCtl or sh*Slc31a1* in plasmids containing a fluorescent Ametrine reporter. Transduced T cells were injected at a 1:1 ratio with shCtl transduced T cells containing a Thy1.1 reporter into *Rag1^-/-^*host mice followed by MOG immunization to induce EAE. Fifteen days after T cell transfer, sh*Slc31a1* transduced T cells were strongly depleted from the spinal cords of host mice compared to shCtl cells (**Fig. 1B**). These results show that CTR1 is critical for the migration, expansion or survival of encephalitogenic T cells in EAE.

**Figure 1:**
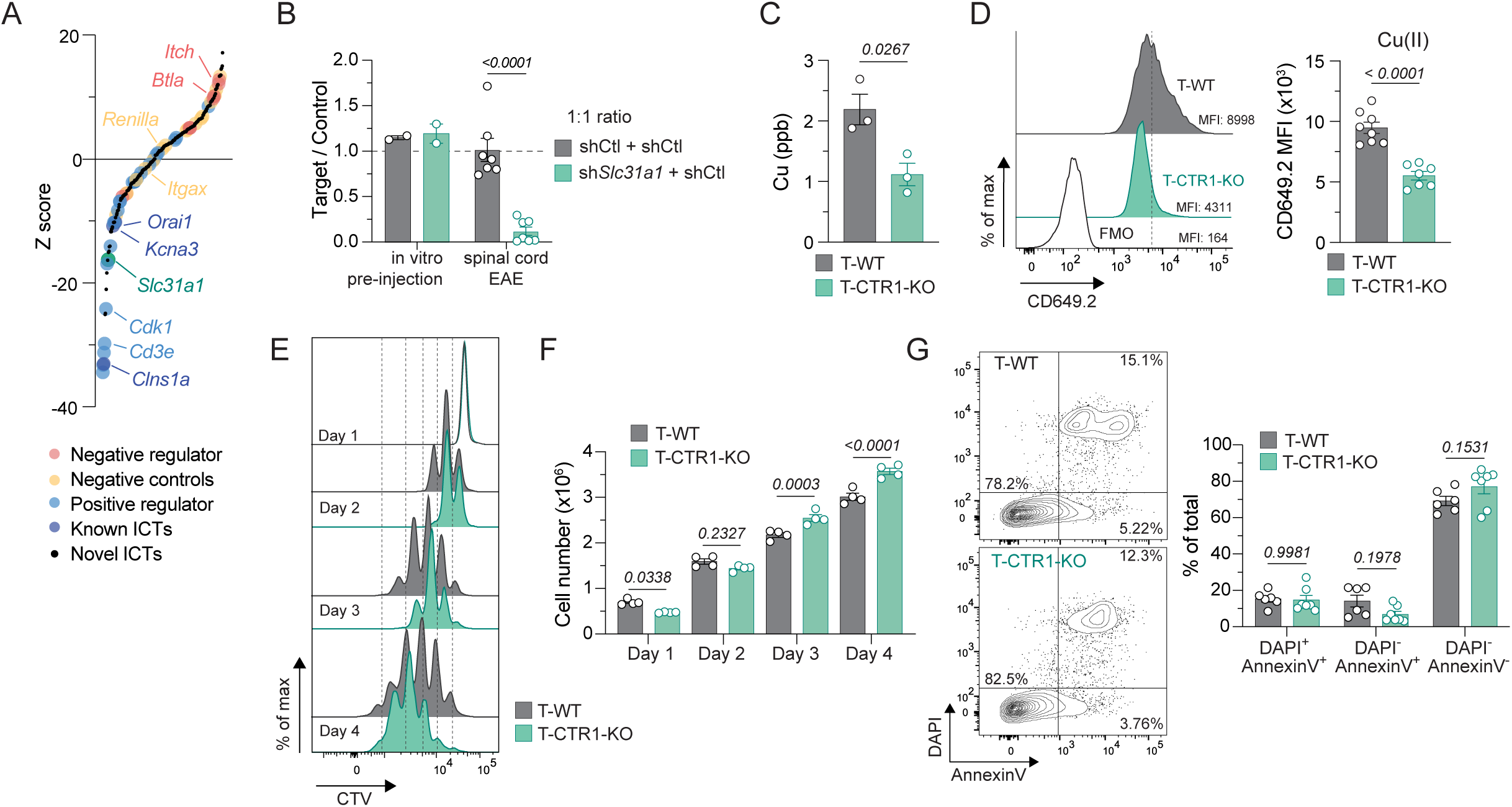
CTR1 (Slc31a1) copper transporter is essential for pT_H_17 cells in EAE. **A**: Z-scores of ICTs and controls in donor 2D2 pT_H_17 cells isolated from the spinal cords of mice with EAE (52 mice pooled from 5 independent screens). **B**: EAE competition assay using shCtrl 2D2 CD4^+^ T cells co-transferred with either shCtrl or sh*Slc31a1* at 1:1 ratio. Quantification of T cell ratios pre-injection and 15 days after MOG immunization (n = 7, 2 independent experiments). **C**: Intracellular copper levels measured in pT_H_17 cells (day 4) from WT mice (T-WT) or T-CTR1-KO mice measured by ICP-MS (n = 3). **D**: Intracellular copper levels in pT_H_17 cells (day 3-5) from WT mice (T-WT) or T-CTR1-KO mice measured by the CD649.2 copper dye (n = 7-8, 2 independent experiments). **E-F**: Proliferation of T-WT and T-CTR1-KO pT_H_17 cells at days 1-4. Representative flow cytometry histograms of Cell Trace Violet (CTV) dilution (**E**). Cell counts (**F**) (n = 4, 2 independent experiments). **G**: Representative flow cytometry plots and quantification of AnnexinV and DAPI positive cells of T-WT and T-CTR1-KO pT_H_17 cells at day 3 (n = 6-7, 2 independent experiments). Statistical significance was calculated by unpaired Student’s *t* test (B-D, F) two-way ANOVA (E, G).

To better understand the role of CTR1 in T cells, we generated mice with T cell specific depletion of CTR1 by crossing *Slc31a1^F/F^*mice with *Cd4Cre* mice, referred to hereafter as T-CTR1-KO mice (and *Slc31a1^F/F^*littermates as T-WT). CD4^+^ T cells isolated from T-CTR1-KO mice had reduced cell surface levels of CTR1 compared to T-WT (**Fig. S1A**). Intracellular copper levels were significantly lower in CD4^+^ T cells from T-CTR1-KO than T-WT mice (**Fig. 1C-D**)^24^, confirming the importance of CTR1 for copper import by T cells. T cell-specific deletion of *Slc31a1* did not lead to gross defects in thymic T cell numbers and thymocyte development (**Fig. S1B-C**). In peripheral lymphoid organs, CD4^+^ T cells numbers were partially reduced in the spleen, but not in LNs (**Fig. S1D**). This was accompanied by a small but significant reduction in the frequencies of effector CD4^+^ T cells (**Fig. S1E**). These data suggest that CTR1 may play a role in CD4^+^ T cells activation and expansion in peripheral lymphoid organs. Moreover, CD8^+^ T cells were markedly reduced in both LNs and spleen (**Fig. S1D**). We also observed a small but significant decrease in CD25^+^FOXP3^+^ regulatory T cell (T_reg_) frequencies in the thymus (**Fig. S1F**), but not in secondary lymphoid organs (**Fig. S1G**), mirroring the findings of a recent study showing no differences in T_reg_ proportions in young mice^23^.

To determine how CTR1 regulates encephalitogenic T cell function, we first investigated the effects of *Slc31a1* deletion in vitro. CTR1-deficient CD4^+^ T cells showed an initial delay in proliferation after anti-CD3/CD28 stimulation, which was followed by increased proliferation, resulting in higher total CD4^+^ T cell counts compared to WT controls 3 and 4 days after activation (**Fig. 1E-F**). Deletion of *Slc31a1* did not affect CD4^+^ T cell viability in vitro (**Fig. 1G**). Moreover, the expression of T cell activation markers CD25, CD44 and CD69 at 24 and 48 hours after anti-CD3/CD28 stimulation was not impaired in CTR1-deficient T cells (**Fig. S2A**). In fact, cell surface levels of CD44 were significantly increased in CTR1-deficient CD4^+^ T cells at 48 hours after stimulation. As CD44 has been proposed to mediate copper import in immune cells^22,23^, this increase may represent a compensatory mechanism to mediate copper transport in the absence of CTR1. However, as intracellular copper is reduced in CTR1-deficient T cells, CD44 upregulation is insufficient to restore intracellular copper levels to those in T-WT CD4^+^ T cells (**Fig. 1C-D**). We conclude that CTR1 is unlikely to regulate encephalitogenic T cells by controlling their activation, viability or proliferation.

### Copper and CTR1 regulate pT_H_17 differentiation and effector function in vitro

Next, we investigated if CTR1 is required for CD4^+^ T cell differentiation by polarizing CD4^+^ T cells from T-CTR1-KO or T-WT mice into different T_H_ and T_reg_ subsets in vitro. In WT T cells, surface CTR1 levels were highest in pT_H_17 cells compared to other CD4^+^ T cell subsets (**Fig. 2A**), suggesting that CTR1 may play a critical role in this T cell subset. *Slc31a1* deletion resulted in a significant lower expression of RORγt (**Fig. 2B**). This defect was specific to pT_H_17 cells (polarized with IL-1β, IL-6, and IL-23), because homT_H_17 cells (polarized with IL-6 and TGFβ) showed no defect in RORγt expression (**Fig. 2C, S2B**). *Slc31a1* deletion did not affect T-bet expression in T_H_1 cells and GATA3 expression in T_H_2 cells (**Fig. 2D, S2C**). Neither did *Slc31a1* deletion impair FOXP3 expression in iT_reg_ cells in vitro (**Fig. 2E, S2D**). Deletion of CTR1 resulted in a strongly impaired production of IL-17A and GM-CSF cytokines in pT_H_17 cells (**Fig. 2F-G, S2E**) and homT_H_17 cells (**Fig. 2H, S2F**), whereas IFN-γ and IL-4 production by T_H_1 and T_H_2 cells, respectively, was unchanged (**Fig. 2I, S2G-H**).

**Figure 2:**
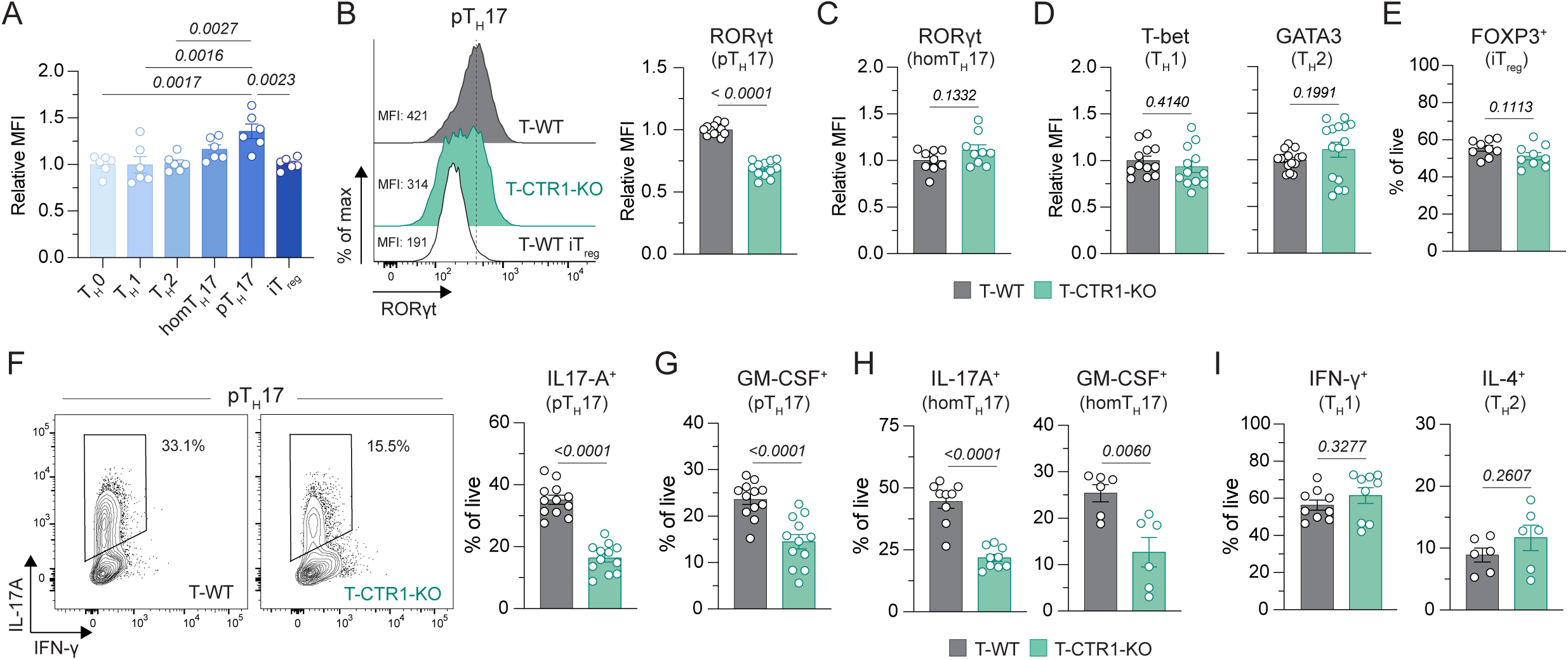
CTR1 regulates pT_H_17 polarization in vitro. **A**: CTR1 expression in CD4 from T-WT and T-CTR1-KO 3 days after activation and polarization in vitro in either T_H_0, T_H_1, T_H_2, homT_H_17, pT_H_17 or iT_reg_ culture conditions, measured by flow cytometry (n = 6, 2 independent experiments). **B**: Representative flow cytometry histogram and quantification of RORγt expression in T-WT and T-CTR1-KO pT_H_17 cells in vitro at day 3, and T-WT iT_reg_ shown as control, relative MFI normalized to T-WT per experiment (n = 12, 4 independent experiments). **C**: RORγt expression in T-WT and T-CTR1-KO homT_H_17 cells in vitro at day 3, relative MFI normalized to T-WT per experiment (n = 9, 3 independent experiments). Representative flow cytometry plot in Fig. S2B. **D**: T-bet (left) and GATA3 (right) expression in T-WT and T-CTR1-KO T_H_1 (left) or T_H_2 (right) cells in vitro at day 3, relative MFI normalized to T-WT per experiment (n = 12-15, 4-5 independent experiments). Representative flow cytometry plot in Fig. S2C. **E**: Frequencies of FOXP3^+^ cells in T-WT and T-CTR1-KO iT_reg_ cells in vitro at day 3 (n = 9, 3 independent experiments). Representative flow cytometry plot in Fig. S2D. **F**: Representative flow cytometry plots and quantification of IL-17A production in T-WT and T-CTR1-KO pT_H_17 cells in vitro at day 3 and after PMA/ionomycin stimulation (n = 12, 5 independent experiments). **G**: GM-CSF production in T-WT and T-CTR1-KO pT_H_17 cells in vitro at day 3 and after PMA/ionomycin stimulation (n = 12, 4 independent experiments). Representative flow cytometry plot in Fig. S2E. **H**: IL-17A and GM-CSF production in T-WT and T-CTR1-KO homT_H_17 cells in vitro at day 3 and after PMA/ionomycin stimulation (n = 9-6, 3-2 independent experiments). Representative flow cytometry plot in Fig. S2F. **I**: IFN-γ (left) and IL-4 (right) production in T-WT and T-CTR1-KO T_H_1 (left) and Th2 (right) cells in vitro at day 3 and after PMA/ionomycin stimulation (n = 9-6, 3-2 independent experiments). Representative flow cytometry plot in Fig. S2G-H. Statistical significance was calculated by one-way ANOVA (A) or unpaired Student’s *t* test (B-I).

To elucidate the role of CTR1 in pT_H_17 differentiation and function, we analyzed global gene expression profiles in T-WT and T-CTR1-KO CD4^+^ T cells polarized into either pT_H_17 or T_H_1 using bulk RNAseq. The number of differentially expressed genes (DEGs) in CTR1-deficient pT_H_17 cells compared to WT pT_H_17 cells was twice as high as the number of DEGs in T_H_1 cells, with 2598 versus 1268 DEGs, respectively (**Fig. S3A-B**). Two thirds of DEGs in T_H_1 cells overlapped with those in pT_H_17 cells, representing a shared dysregulated transcriptional program in the absence of CTR1 (**Fig. S3C**). Pathway and upstream regulator analyses of DEGs revealed that both CD4^+^ T cell subsets share the upregulation of pathways related to cell proliferation and cell cycle (**Fig. S3D-E**), consistent with the robust cell expansion observed in vitro (**Fig. 1F**). Both subsets also showed a pronounced dysregulation of cytokine signaling, with a notable downregulation of STAT3, IL-1β and TGFβ, upstream regulators essential for T_H_17 polarization (**Fig. S3E**). In addition, network clustering analysis of pT_H_17 cells showed a downregulation of cytokine signaling and inflammation pathways despite the pronounced upregulation of immune cell activation and cell cycle pathways (**Fig. S3F**). Importantly, transcriptomic analyses of DEGs revealed that CTR1-deficient pT_H_17 cells failed to upregulate many key T_H_17 signature genes (e.g. *Il17a/f, Ccl5, Il23r*), while T_H_1 signature genes were not affected by the deletion of *Slc31a1*, confirming a specific differentiation defect in pT_H_17 cells (**Fig. 3A**). This finding is further corroborated by gene set enrichment analysis (GSEA) showing significant depletion of genes upregulated in pT_H_17 cells along with the enrichment of genes downregulated in homT_H_17 cells (**Fig. 3B**). Together, our findings demonstrate that CTR1 is essential to establish the pT_H_17 transcriptional program, but not the T_H_1 program.

**Figure 3:**
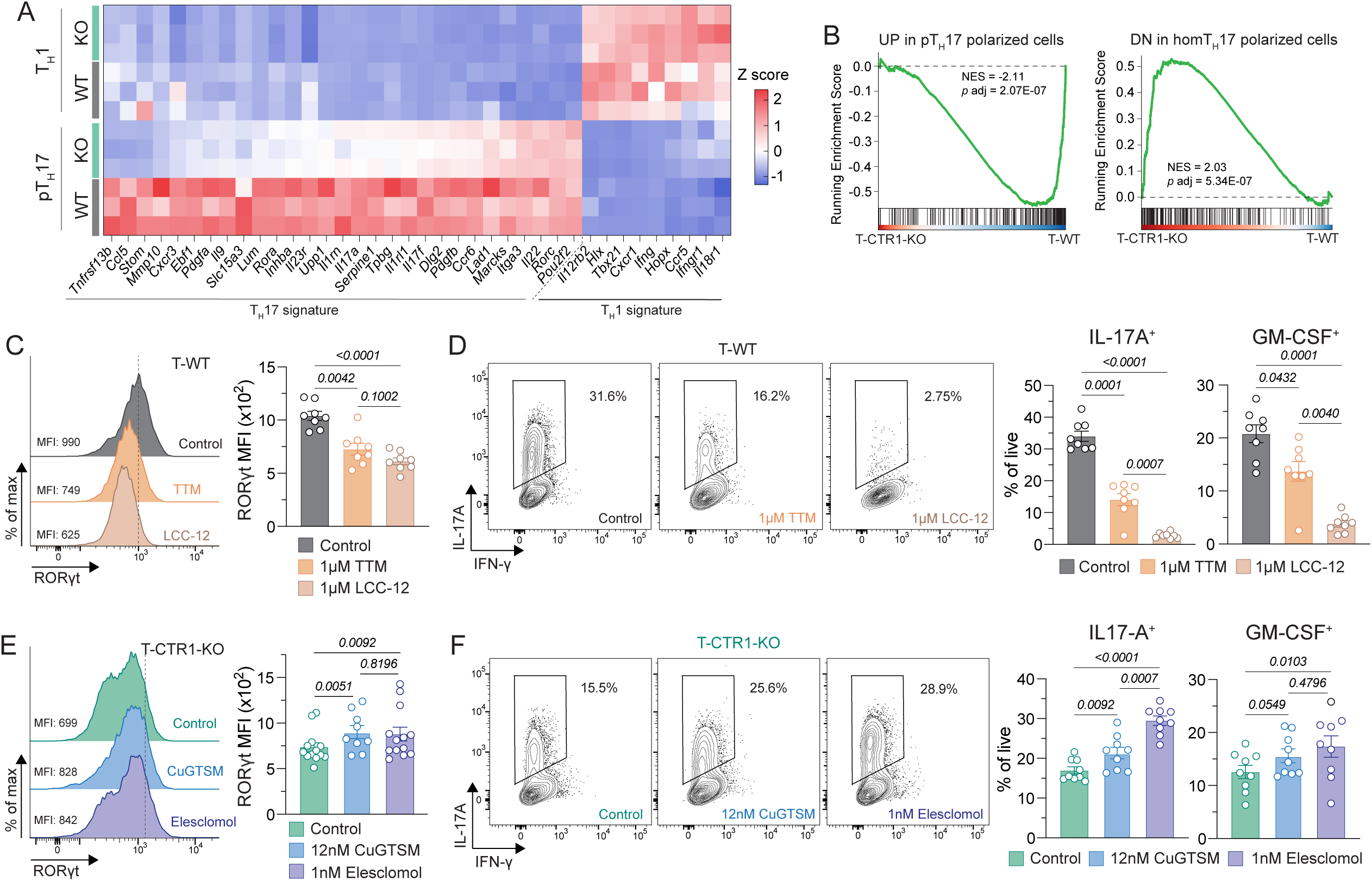
Copper and CTR1 are essential to pT_H_17 polarization and function in vitro. **A**: Bulk-RNAseq analysis of T-CTR1-KO versus T-WT CD4^+^ T cells cultured in either pT_H_17 or T_H_1 conditions in vitro for 3 days, heatmap showing key pTh17 and Th1 signature genes (n = 3). **B**: GSEA of T-CTR1-KO versus T-WT pT_H_17 cells; “pT_H_17 cells DOWN” GSE39820; “homT_H_17 cells UP” GSE21670. **C-D**: T-WT pT_H_17 cells treated with 1 µM tetrathiomolybdate (TTM), 1 µM LCC-12 or vehicle control for 3 days. Representative flow cytometry histogram and quantification of RORγt expression (**C**) (n = 8, 2 independent experiments). Representative flow cytometry plots of IL-17A and quantification of IL-17A and GM-CSF production after PMA/ionomycin stimulation (**D**) (n = 8, 2 independent experiments). **E-F**: T-CTR1-KO pT_H_17 cells treated with 12 nM CuGTSM, 1nM Elesclomol or control for 3 days. Representative flow cytometry histogram and quantification of RORγt expression (**E**) (n = 13-9, 4-3 independent experiments). Representative flow cytometry plots of IL-17A and quantification of IL-17A and GM-CSF production after PMA/ionomycin stimulation (**F**) (n = 9, 3 independent experiments). Statistical significance was calculated by one-way ANOVA (C-D, E-F), mixed-effects analysis (E).

To validate the importance of copper for pT_H_17 polarization and function, we altered intracellular copper levels during pT_H_17 differentiation in vitro. First, we polarized T-WT CD4^+^ T cells into pT_H_17 cells in the presence of the copper chelators tetrathiomolybdate (TTM) or LCC-12. Copper chelation significantly reduced transcript levels of T_H_17 signature genes including *Rora* and *Inhba* (encoding an Activin-A subunit) compared to vehicle control treatment (**Fig. S3G**). At the protein level, we found significantly impaired expression of RORγt (**Fig. 3C**) and production of IL-17A and GM-CSF in copper chelator-treated T-WT cells (**Fig. 3D**). Importantly, treatment of T-CTR1-KO cells with the cell-permeable copper complex CuGTSM or the copper ionophore Elesclomol rescued the defects in pT_H_17 polarization and function, resulting in significantly increased levels of T_H_17 transcripts (**Fig. S3H**), RORγt expression (**Fig. 3E**) and IL-17A and GM-CSF production (**Fig. 3F**) compared to vehicle control treated cells, demonstrating that CTR1 regulates pT_H_17 cell function through copper import. Taken together, our data show that copper and CTR1 are critical for the differentiation and function of pT_H_17 in vitro, but largely dispensable for other CD4^+^ T cell subsets.

### Copper and CTR1 shape T cell metabolism to regulate pT_H_17 differentiation

We next investigated the mechanisms by which copper and CTR1 regulate pT_H_17 polarization. Copper has essential functions in COX assembly and electron transfer in mitochondrial respiration^16,23^. Consistent with this role of copper, T-CTR1-KO pT_H_17 cells had lower levels of mitochondrial reactive oxygen species (mROS) compared to T-WT cells (**Fig. 4A**). We also observed a decreased basal oxygen consumption rate (OCR) and decreased maximal OCR capacity, readouts of oxidative phosphorylation (OXPHOS) by mitochondria (**Fig. 4B**). The reduced OCR was not specific to pT_H_17 cells but also present in CTR1-deficient T_H_1 cells (**Fig. S4A-B**). The defect in mitochondrial respiration was not secondary to T cell differentiation, since we observed similar defects already at 24 hours after CD4^+^ T cell stimulation at the beginning of T_H_1 or pT_H_17 polarization (**Fig. S4C-D**). To understand the metabolic consequences of impaired mitochondrial respiration, we analyzed metabolites in WT and CTR1-deficient pT_H_17 cells by liquid chromatography–mass spectrometry (LC-MS). The levels of dihydroorotate, glycerol-3-phosphate (G3P) and malate were strongly increased in T-CTR1-KO cells compared to controls, consistent with impaired ETC function and reduced activity of G3P and malate shuttles (**Fig. 4C, S4E**). This conclusion is further supported by a decreased NAD^+^/NADH ratio in pT_H_17 cells from T-CTR1-KO compared to T-WT mice (**Fig. 4D**). In parallel, CTR1-deficient pT_H_17 cells showed increased baseline glycolysis as indicated by a higher extracellular acidification rate (ECAR) and lactate accumulation (**Fig. 4E-F**). Indeed, T-CTR1-KO pT_H_17 cells become highly dependent on glucose to produce ATP (**Fig. S4F-G**) which likely supports their protein production and sustains their proliferative capacity. Moreover, CTR1-deficient pT_H_17 cells show enhanced glucose usage through glycolysis, a potential increase in polyol pathway activity as suggested by the increased expression of *Sord* (sorbitol dehydrogenase) (**Fig. S4H**) and increased usage of the pentose phosphate shunt as indicated by the higher sedoheptulose-7-phosphate levels (**Fig. 4F**). As SORD utilizes NAD^+^, its increased expression may contribute to the lower NAD^+^/NADH ratio in CTR1-deficient pT_H_17 cells. Our data indicate that impaired mitochondrial respiration in CTR1-deficient pT_H_17 cells results in a multi-pathway metabolic overhaul consisting of augmented glycolysis, impaired mitochondrial TCA cycle, and active pyrimidine and purine synthesis.

**Figure 4:**
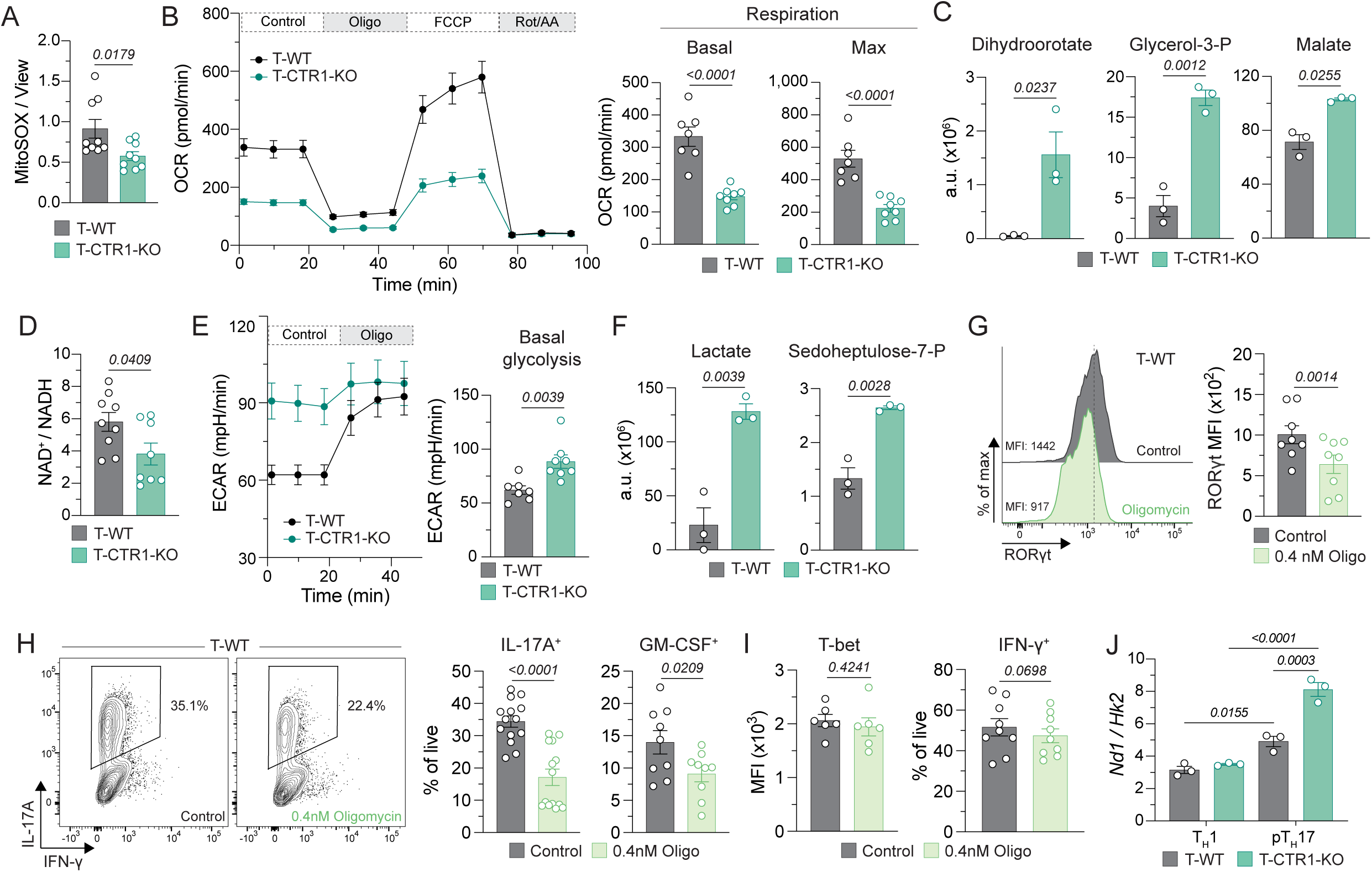
CTR1 regulates pT_H_17 via T cell metabolism. **A**: ROS production in mitochondria measured by MitoSOX normalized over MitoView, in T-WT and T-CTR1-KO pT_H_17 cells day 3 (n = 9, 3 independent experiments). **B**: Time course and quantification of OCR (oxygen consumption rate) T-WT and T-CTR1-KO pT_H_17 cells at day 3. Basal respiration is quantified as a mean of 0-20 min, Max respiration is quantified as a mean of 50-70min (n = 7-8, 3 independent experiments). **C**: Dihydroorotate, glycerol-3-P and malate levels in T-WT and T-CTR1-KO pT_H_17 cells day 3 measured by LC-MS (n = 3). **D**: NAD^+^ / NADH ratio in T-WT and T-CTR1-KO pT_H_17 cells at day 3 (n = 8-9, 2 independent experiments). **E**: Time course and quantification of ECAR (extracellular acidification rate) in T-WT and T-CTR1-KO pT_H_17 cells at day 3. Basal glycolysis is quantified as a mean of 0-20 min (n = 7, 3 independent experiments). **F**: Lactate and sedoheptulose-7-P levels in T-WT and T-CTR1-KO pT_H_17 cells day 3 measured by LC-MS (n = 3). **G-H**: T-WT pT_H_17 cells treated with 0.4 nM oligomycin or vehicle control for 3 days. Representative flow cytometry histogram and quantification of RORγt expression (**G**) (n = 8, 3 independent experiments). Representative flow cytometry plots of IL-17A and quantification of IL-17A and GM-CSF production after PMA/ionomycin stimulation (**H**) (n = 14-9, 5-3 independent experiments). **I**: T-bet expression and IFN-γ production in T-WT T_H_1 cells treated with 0.4 nM oligomycin or vehicle control for 3 days (n = 6-9, 2-3 independent experiments). Representative flow cytometry plot in Fig. S4J. **J**: Mitochondria copy number measured by *Nd1:Hk2* ratio from bulk RNAseq (n = 3). Statistical significance was calculated by unpaired Student’s *t* test (A-F), paired Student’s *t* test and one-way ANOVA (J).

Mitochondrial respiration is essential for T_H_17 polarization^5,25–29^. We found that prolonged treatment of T-WT CD4^+^ T cells with 0.4 nM of oligomycin, an ATP synthase (ETC complex V) inhibitor, for 3 days during pT_H_17 polarization reduced their OCR to levels comparable to those in T-CTR1-KO cells (**Fig. S4I**). Moreover, pT_H_17 cell differentiation and function were decreased in oligomycin treated cells as evidenced by lower RORγt levels and decreased IL-17A and GM-CSF production (**Fig. 4G-H**). By contrast, T_H_1 cells were not affected by oligomycin treatment because T-bet expression and IFN-γ production were comparable to vehicle control-treated cells (**Fig. 4I, Fig. S4J**). Of note, mitochondrial copy numbers were strongly increased in pT_H_17 cells from T-CTR1-KO mice compared to either T-WT pT_H_17 cells or CTR1-deficient T_H_1 cells (**Fig. 4J**), potentially as a compensatory mechanism for their impaired respiratory function. Taken together, our data show that CTR1 and copper shape T cell metabolism and promote pT_H_17 polarization and function in vitro by sustaining mitochondrial respiration.

### Copper and CTR1 do not regulate pT_H_17 cells via SOD1, ULK1/2 or MEK1/2

Besides regulating COX assembly and mitochondrial respiration, copper is a cofactor for other enzymes involved in various cellular processes. Superoxide dismutase 1 (SOD1), which is located in the cytosol and mitochondrial intermembrane space, requires both copper and zinc to reduce reactive oxygen species (ROS). Controlling ROS levels in T cells is essential to balance the need for ROS-mediated T cell activation with ROS toxicity and inhibitory effects on metabolic reprogramming^30^. We hypothesized that reduced cytosolic copper levels in T-CTR1-KO CD4^+^ T cells result in impaired SOD1 function and increased ROS levels. T-CTR1-KO CD4^+^ T cells had reduced SOD activity compared to T-WT cells (**Fig. S5A**), resulting in cytosolic ROS accumulation in both T_H_1 and pT_H_17 cells **(Fig. S5B**). To evaluate the importance of SOD1 in T_H_17 polarization and function in vitro, we treated T-WT CD4^+^ T cells with the SOD1 inhibitor LCS-1 during pT_H_17 differentiation. Prolonged SOD1 inhibition did not significantly affect RORγt expression (**Fig. S5C**) or T_H_17 cytokine production (**Fig. S5D**). To confirm that CTR1 does not regulate T_H_17 polarization through SOD1 activity, we treated T-CTR1-KO CD4^+^ T cells with the ROS scavenger Tiron during T cells activation and pT_H_17 polarization. ROS scavenging in T-CTR1-KO cells did not rescue RORγt expression (**Fig. S5E**) or T_H_17 cytokine production (**Fig. S5F**). Together these findings indicate that CTR1 does not control T_H_17 polarization and function via SOD1 activity.

Copper was also reported to interact with UNC-51-like kinases 1 and 2 (ULK1/2), thereby regulating the formation of autophagosomes^31^. Autophagy is involved in inflammation and T_H_17 polarization and function^32^. We first measured autophagy levels in T-CTR1-KO and T-WT pT_H_17 cells via LC3-II formation but did not find significant differences in autophagy levels between WT and CTR1-deficient pT_H_17 cells (**Fig. S5G**). The autophagy inhibitor chloroquine (CQ) was reported to prevent T_H_17 polarization and to protect mice from EAE^32^. Because the QC concentrations used in this study impaired T cell activation, we treated T-WT CD4^+^ pT_H_17 cells with a low dose of CQ that did not affect T cell activation or viability but failed to observe effects of autophagy inhibition on RORγt expression (**Fig. S5H**) and T_H_17 cytokine production in vitro (**Fig. S5I**). We conclude that CTR1-deficient pT_H_17 cells do not have a defect in autophagy, and that copper does not regulate pT_H_17 cell function through autophagy.

The MAPK/ERK kinases 1 and 2 (MEK1/2) bind copper to sustain their activity^33,34^. ERK1/2, the targets of MEK1/2 kinases, have been linked to T_H_17 cell function but with conflicting results. While one study showed that ERK2 phosphorylates RORγt and inhibits its function^35^, another reported that ERK inhibition decreases T_H_17 cells favoring T_reg_ cells in colitis^36^. To explore if deletion of *Slc31a1* influences MEK1/2 function in pT_H_17 cells, we measured ERK phosphorylation in naïve T-CTR1-KO and T-WT cells. We did not observe differences in phospho-ERK levels in WT and CTR1-KO cells before or 10 minutes after anti-CD3/CD28 stimulation (**Fig. S5J**). Taken together, our data indicate that CTR1 does not promote T_H_17 polarization by regulating SOD1 function, autophagy or MAPK signaling, reinforcing the role of copper in mitochondrial respiration to control T_H_17 cells.

### CTR1 regulates DNA methylation of pT_H_17 signature genes

CTR1 deletion profoundly reshapes T cell metabolism and the metabolite landscape (**Fig. S4E**). Notably, T-CTR1-KO pT_H_17 cells showed a marked increase in the ratio of 2-hydroxyglutarate (2-HG), succinate, and fumarate relative to α-ketoglutarate (αKG) (**Fig. 5A**). This metabolic shift is relevant to epigenetic regulation, as 2-HG, succinate, and fumarate are potent inhibitors of TET and histone demethylase (HDM) enzymes responsible for the removal of methylation marks on DNA and histones, respectively, while αKG serves as the essential co-substrate required for their enzymatic activity^37,38^. Moreover, treatment of T_H_17 cells with 2-HG has been shown to decreased IL-17A production^28^. We hypothesized that the elevated ratios of 2-HG/αKG, succinate/αKG, and fumarate/αKG observed in T-CTR1-KO pT_H_17 cells are associated with inhibition of DNA demethylating enzymes and thus, cause the dysregulation of pT_H_17 differentiation^39^. To test this hypothesis, we polarized T-WT CD4^+^ T cells into pT_H_17 cells in the presence of 2-HG, succinate or fumarate. Treatment of T cells with 200 µM 2-HG for 3 days during pT_H_17 polarization resulted in significantly lower in RORγf expression (**Fig. 5B**) and IL-17A production (**Fig. 5C**) compared to vehicle control. Similarly, treatment with 100 µM succinate resulted in lower RORγt and IL-17A expression, whereas 200 µM fumarate only reduced IL-17A production (**Fig. S6A, B**). By contrast, treatment of T-WT cells with 200 µM 2-HG during T_H_1 differentiation did not affect T-bet expression and IFN-γ production (**Fig. 5D, S6C**). We conclude that the elevated ratios of 2-HG, succinate, and fumarate relative to αKG in CTR1-deficient cells suppress pT_H_17 differentiation and function.

**Figure 5:**
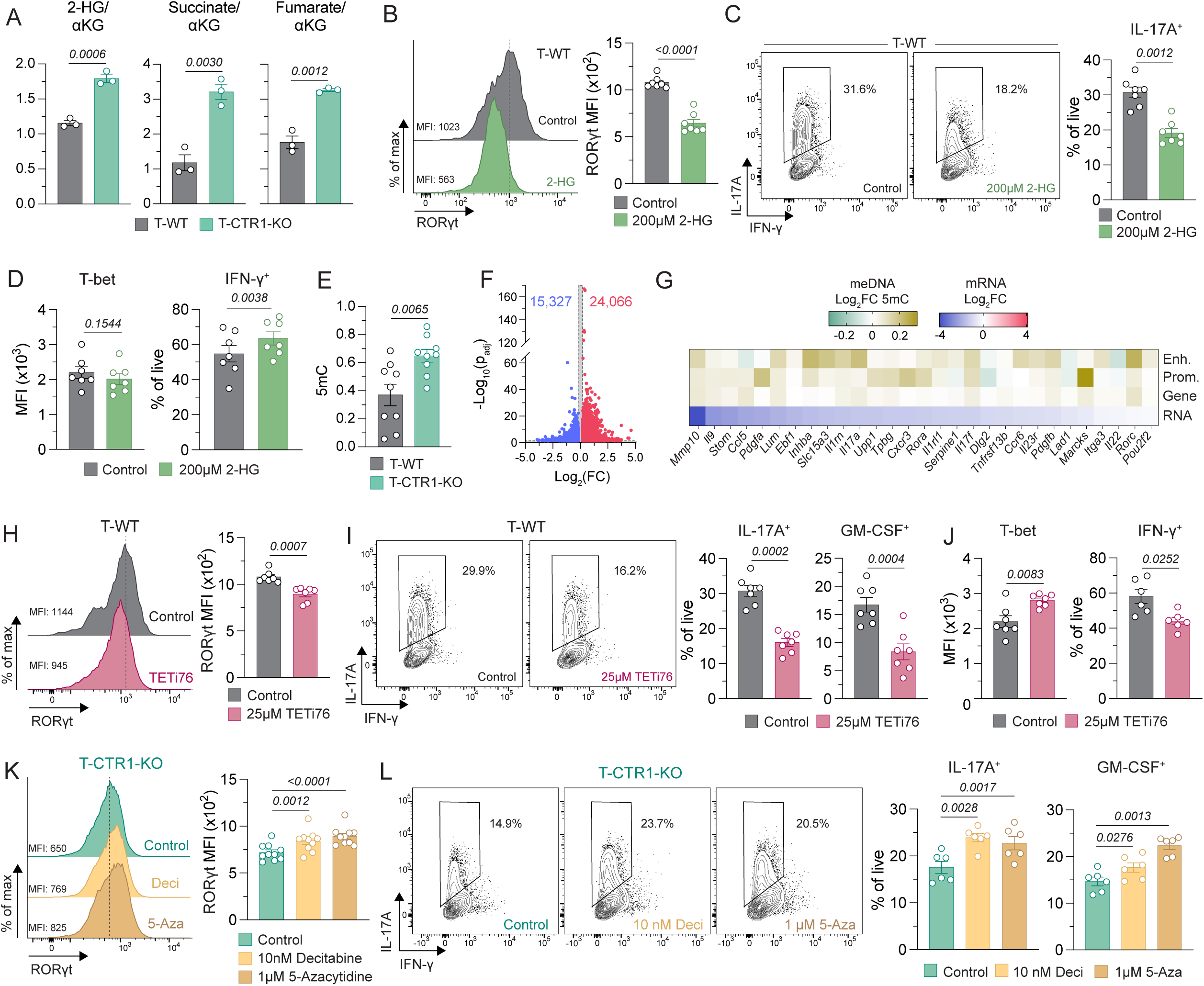
Copper and CTR1 regulate DNA methylation in T cells. **A**: Ratios of 2-hydroxyglutarate (2-HG), succinate and fumarate and levels over α-ketoglutarate (α-KG) in T-WT and T-CTR1-KO pT_H_17 cells at day 3 measured by LC-MS (n = 3). **B-C**: T-WT pT_H_17 cells treated with 200µM 2-HG or vehicle control for 3 days. Representative flow cytometry histogram and quantification of RORγt expression (**B**) (n = 7, 2 independent experiments). Representative flow cytometry plots and quantification of IL-17A production after PMA/ionomycin stimulation (**C**) (n = 7, 2 independent experiments). **D**: T-bet expression and IFN-γ production in T-WT T_H_1 cells treated with 200 µM 2-HG or vehicle control for 3 days. Representative flow cytometry plot in Fig. S6C. **E**: Global 5mC DNA methylation levels in T-WT and T-CTR1-KO pT_H_17 cells at day 3 measured by ELISA (n = 9, 2 independent experiments). **F-G**: 5mC DNA methylation sequencing in T-WT and T-CTR1-KO pT_H_17 cells at day 3 (n = 2). **F**: Volcano plot of differentially methylated regions (1 kb windows) (n = 2). **G**: Heatmap of T_H_17 signature genes showing differential 5mC for at enhancer, promoter, whole gene level (n = 2), and mRNA expression (bulk RNAseq as shown in Figure 3A, n=3). **H-I**: T-WT pT_H_17 cells treated with 25µM TETi76 or vehicle control for 3 days. Representative flow cytometry histogram and quantification of RORγt expression (**H**) (n = 7, 2 independent experiments). Representative flow cytometry plots of IL-17A and quantification of IL-17A and GM-CSF production after PMA/ionomycin stimulation (**I**) (n = 7, 2 independent experiments). **J**: T-bet expression and IFN-γ production in T-WT T_H_1 cells treated with 25 µM TETi76 or vehicle control for 3 days (n = 7-6, 2 independent experiments). Representative flow cytometry plot in Fig. S6G. **K-L**: T-WT pT_H_17 cells treated with 10 nM decitabine, 1 µM 5-azacytidine or vehicle control for 3 days. Representative flow cytometry histogram and quantification of RORγt expression (**K**) (n = 10, 3 independent experiments). Representative flow cytometry plots of IL-17A and quantification of IL-17A and GM-CSF production after PMA/ionomycin stimulation (**L**) (n = 6, 2 independent experiments). Statistical significance was calculated by unpaired Student’s *t* test (A, E), paired Student’s *t* test (B-D, H-J) or one-way ANOVA (K-L).

We next analyzed global DNA methylation levels by measuring 5-methylcytosine (5mC)-modified DNA which was significantly increased in CTR1-deficient pTh17 cells compared to WT controls (**Fig. 5E**). DNA methylation sequencing of pT_H_17 cells revealed that T-CTR1-KO cells had 24,066 hypermethylated regions compared to T-WT cells (**Fig. 5F**). Focusing on select T_H_17 signature genes, we observed increased DNA methylation of enhancers, promoters or whole gene regions (**Fig. 5G**). The hypermethylation of entire gene regions and promoters showed a moderate correlation with decreased transcript levels of T_H_17 genes in T-CTR1-KO pT_H_17 cells, whereas no correlation was observed between hypermethylation of enhancer regions and transcript levels (**Fig. S6D**). Hypermethylation at enhancer regions did, however, correlate with decreased chromatin accessibility in T-CTR1-KO pT_H_17 cells as measured by ATAC-sequencing (**Fig. S6E**). Taken together, our results demonstrate that CTR1-deficient pT_H_17 cells have hypermethylated DNA in regions that regulate the expression of signature T_H_17 genes, providing a link between metabolic rewiring, epigenetic regulation and expression of genes associated with T_H_17 differentiation.

We evaluated the significance of DNA methylation for pT_H_17 differentiation and function. First, we cultured T-WT CD4^+^ T cells under pT_H_17 polarizing conditions in the presence of 25 µM TETi76, a TET inhibitor. Inhibition of TET enzymes impaired pT_H_17 cell differentiation and function, as apparent from the decreased expression of T_H_17 signature genes (**Fig. S6F**), RORγt (**Fig. 5H**), and IL-17A and GM-CSF (**Fig. 5I**) compared to vehicle control. By contrast, treatment of T-WT cells with TETi76 during T_H_1 polarization resulted in higher T-bet expression and lower IFN-γ production (**Fig. 5J, Fig. S6G**). Next, we treated CTR1-deficient T cells with DNA methyltransferase (DNMT) inhibitors during pT_H_17 differentiation. Culture of T-CTR1-KO cells with either decitabine or 5-azacytidine resulted in significantly higher expression of RORγt (**Fig. 5K**), as well as IL-17A and GM-CSF (**Fig. 5L**) compared to CTR1-deficient cells treated with vehicle control. Moreover, 5-azacytidine treatment increased the expression of pT_H_17 signature genes in T-CTR1-KO T cells (**Fig. S6H**). Collectively, these data demonstrate that DNA demethylation is dependent on CTR1 and is essential for the differentiation and function of T_H_17 cells but has little effect on T_H_1 cells.

DNA methylation is dynamic and can modulate transcription via several mechanisms,^40^ including the recruitment of histone deacetylases (HDACs). Once recruited HDACs can repress transcription by removing activating histone marks such as histone 3 lysine 27 acetylation (H3K27Ac). A recent study demonstrated that *Slc31a1* deletion results in lower H3K27Ac levels in T_reg_ cells^23^. Moreover, HDACs were shown to modulate homT_H_17 polarization^41,42^. We hypothesized that hypermethylated DNA in T-CTR1-KO results in increased recruitment of HDACs, decreased H3K27Ac and impaired pT_H_17 differentiation. To test this hypothesis, we differentiated T-WT CD4^+^ T cells into pT_H_17 cells in the presence of the histone acetyl transferase (HAT) inhibitors anacardic acid, CPTH2 or A-485. Inhibition of HATs did not affect RORγt expression but significantly reduced IL-17A production compared to vehicle control treated cells (**Fig. S6I**). Next, we treated T-CTR1-KO CD4^+^ T cells with HDAC inhibitors to prevent H3K27Ac removal. Trichostatin A (TSA) or SAHA treatment did not increase RORγt levels compared to vehicle control treated cells (**Fig. S6J, K**) but significantly increased IL-17A production by T-CTR1-KO cells. Collectively these data indicate that hypermethylation of CTR1-deficient T cells may inhibit pT_H_17 function by interfering with activating histone acetylation marks.

### CTR1 regulates chromatin accessibility and the recruitment of AP-1 to T_H_17 signature genes

We hypothesized that DNA hypermethylation of CTR1-deficient T cells alters their chromatin landscape. Therefore, we next explored the effects of CTR1 deletion on chromatin accessibility in pT_H_17 cells using ATAC-sequencing. *Slc31a1* deletion resulted in profound changes in chromatin accessibility in pT_H_17 cells with the closing of 134,559 regions and the opening of 17,569 regions (**Fig. 6A**). We leveraged these data to predict transcription factor (TF) binding using several motif analysis platforms. All approaches showed a significant depletion of members of the AP-1 (activating protein 1) family of TFs in CTR1-deficient pT_H_17 cells compared to WT control pT_H_17 cells (**Fig. 6B-D, S7A, B**). AP-1 family TFs, which contain a bZIP (basic leucine zipper) motif that allows them to homo- or heteromultimerize, play an essential role in pT_H_17 specification. BATF (bZIP activating transcription factor) is a pioneering TF for T_H_17 cells^43,44^, dimerizing with other AP-1 members such as the JunB and Fosl2 dimer, the top enriched motif in the HOMER *de novo* analysis. BATF can also cooperate with IRF4 (interferon regulating factor 4) to initiate the T_H_17 program^45,46^. The reduced binding of BATF, Fosl2 and JunB in CTR1-deficient pT_H_17 cells predicted by motif analysis is consistent with the impaired expression of the pT_H_17 genes (**Fig. 3A, B**). Moreover, we observed the depletion of binding motifs for STAT3 and RORγt TFs, which enforce pT_H_17 differentiation, in T-CTR1-KO pT_H_17 cells (**Fig. 6B, C**). A TF occupancy analysis using TOBIAS also predicted the enrichment of E2F family members in T-CTR1-KO pT_H_17 cells (**Fig. 6B**), which may explain the upregulation of cell cycle pathways we observed (**Fig. S4D, F**). A TF binding motif analysis using HOMER predicted the enrichment of ETS family transcription factors in CTR1-deficient pT_H_17 cells (**Fig. 6D, S7B**). Ets1 is a negative regulator of Th17 differentiation and represses RORγt and IL-17A expression^47^. It is also required for the recruitment of CTCF to poise the chromatin for T_H_ specification^43^, pointing to a potential role of CTR1 in the regulation of 3D chromatin structure.

**Figure 6:**
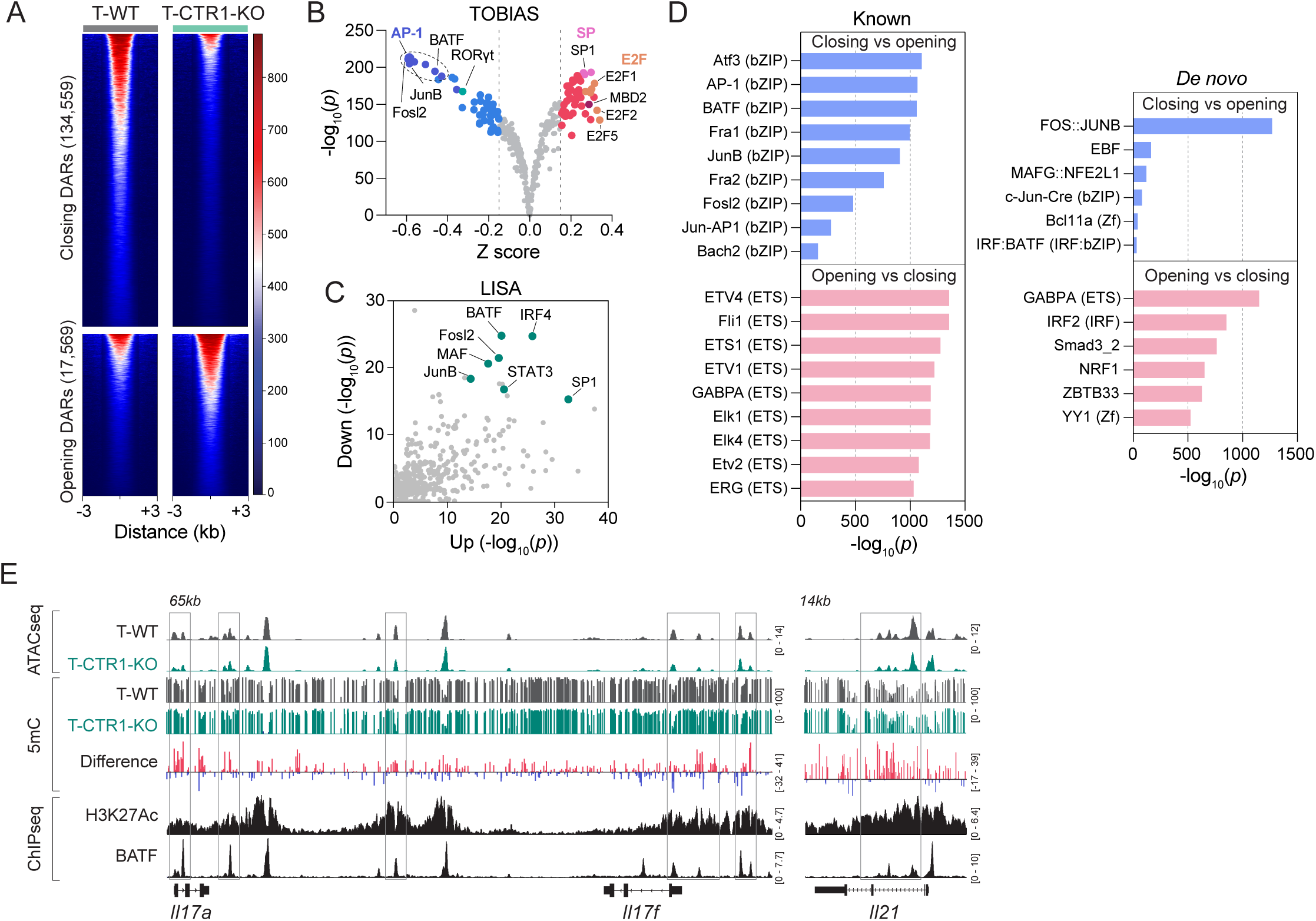
Copper and CTR1 regulate chromatin opening and AP-1 transcription factor recruitment to key T_H_17 loci. **A**: Global differential chromatin opening and closing regions in T-CTR1-KO versus T-WT pT_H_17 cells at day 3 measured by ATACseq (n = 3). **B**: Z scores and significance of enriched and depleted transcription factors in T-CTR1-KO versus T-WT pT_H_17 cells shown in (A) analyzed by TOBIAS. **C**: Predicted dysregulated transcription factors in T-CTR1-KO versus T-WT pT_H_17 cells shown in (A) analyzed by LISA. **D**: Top dysregulated transcription factors in T-CTR1-KO versus T-WT pT_H_17 cells shown in (A) analyzed by HOMER. **E**: Example traces for *Il17a-f* and *Il21* loci showing representative ATACseq and 5mC of T-WT and T-CTR1-KO pT_H_17 cells at day 3, published ChIPseq binding profiles for H3K27Ac (GSM2905773) and BATF (GSM1004796) in T_H_17 cells.

DNA methylation was previously reported to regulate the transcription of the *Il17a/f* locus^48,49^, which is essential for T_H_17 specification. We found that T-CTR1-KO pT_H_17 cells have increased 5mC DNA methylation at several key regions of the *Il17a/f* locus that correlate with BATF binding and H3K27Ac marks of active transcription (**Fig. 6E**). Some of these regions were also associated with decreased chromatin accessibility in the ATAC-seq data (**Fig. 6E**). We identified similar patterns of DNA hypermethylation in other key T_H_17 genes (**Fig. 5G, S6D-E**) including *Il21*, a cytokine produced by CD4^+^ T cells that promotes T_H_17 differentiation (**Fig. 6E**). Together, our data demonstrate that CTR1-deficient T cells have a profoundly altered chromatin accessibility landscape that is associated with reduced recruitment of AP-1 family TFs known to pioneer T_H_17 differentiation. Importantly, key regulatory regions of T_H_17 genes such as *Il17a*, *Il17f* and *Il21*, show increased DNA methylation at H3K27Ac-marked transcriptionally active regions bound by BATF, providing a link between impaired pT_H_17 differentiation and copper depletion.

### Copper and CTR1 are essential for the pathogenicity of T_H_17 cells in EAE

Given the importance of CTR1 for pT_H_17 differentiation in vitro, we investigated its role in the encephalitogenic function of pT_H_17 cells in EAE. Of note, *Slc31a1* expression was significantly elevated in CD4^+^ T cells infiltrating the spinal cords of WT mice with EAE compared to CD4^+^ T cells isolated from their spleen (**Fig. 7A**), suggesting a higher expression of CTR1 in autoreactive T cells. By comparison, the mRNA levels of *Slc31a2* (CTR2), *Atp7a* (encoding a copper-transporting P-type ATPase mutated in Menkes disease) and *Cd44* (which has been associated with copper regulation^22,23,50^) were unchanged (**Fig. S8A**). Significantly elevated *SLC31A1* mRNA expression was also observed in circulating effector T cells (T_eff_) isolated from the blood of patients with relapsing remitting MS (RRMS) compared to healthy controls (HC) (**Fig. 7B**). The expression of *SLC31A2* and *ATP7A* was unchanged, whereas *CD44* mRNA levels were significantly elevated in patients (**Fig. S8B**). Interestingly, the high expression of *Slc31a1* in T cells infiltrating spinal cords of mice with EAE and T_eff_ of patients with RRMS was accompanied by increased mRNA levels of *Slc25a3* and *Cox17*, which have both been implicated in copper delivery pathways that support COX assembly in mitochondria (**Fig. 7C, D**)^51^. The high expression of proteins transporting copper across the PM and into mitochondria suggests a dependency of pT_H_17 cells on copper to sustain their encephalitogenic function.

**Figure 7:**
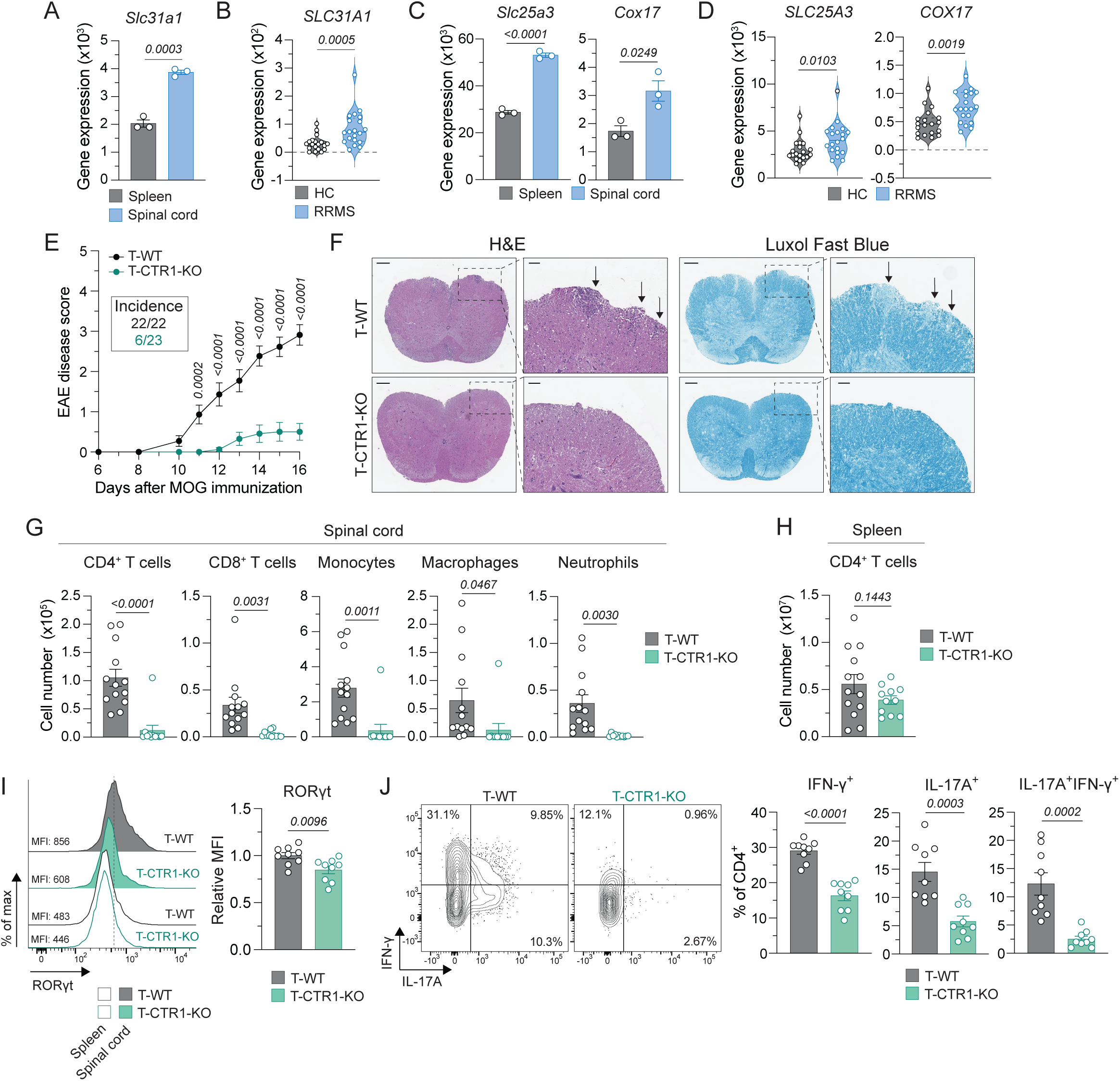
CTR1 is essential for the pathogenicity of T_H_17 cells in EAE. **A**: *Slc31a1* mRNA expression in donor 2D2 pT_H_17 cells isolated from spleen and spinal cords of mice with EAE, measured by bulk RNAseq (n = 3). **B**: *SLC31A1* mRNA expression in blood T_eff_ from healthy control (HC) and patients with active relapsing-remitting multiple sclerosis (RRMS) measured by bulk RNAseq. **C**: *Slc25a3* and Cox17 mRNA expression in donor 2D2 pT_H_17 cells isolated from spleen and spinal cords of mice with EAE as shown in (A). **D**: *SLC25A3* and *COX17* mRNA expression in blood T_eff_ from HC and RRMS patients as shown in (B). **E-J**: Active EAE in T-WT and T-CTR1-KO mice. **E**: EAE disease score and incidence (n = 22-23, 3 independent experiments). **F**: Representative histology of spinal cords with H&E and Luxol Fast Blue staining. Scale bars represent 200µm and 50µm. **G**: Number of immune cells in spinal cords (n = 13, 2 independent experiments). **H**: Number of CD4^+^ T cells in spleen (n = 13, 2 independent experiments). **I**: Representative flow cytometry histogram and quantification of RORγt expression in T cells isolated from spinal cords and spleen shown as control, relative MFI normalized to T-WT per experiment (n = 9, 2 independent experiments). **J**: Representative flow cytometry plots and quantification of IL-17A and IFN-γ production in T cells isolated from spinal cord and restimulated with PMA/ionomycin (n = 9, 2 independent experiments). Statistical significance was calculated by unpaired Student’s *t* test (A, C, I-J), Mann-Whitney test (B, D), two-way ANOVA (E) or Welch’s *t* test (G, H).

To test this hypothesis, we induced active EAE in T-WT and T-CTR1-KO mice. Mice lacking CTR1 in T cells were strongly protected from developing EAE with only 6 of 23 T-CTR1-KO mice developing the disease and exhibiting only mild disease symptoms compared to T-WT mice (**Fig. 7E**). Spinal cords of T-CTR1-KO mice showed no evidence of inflammation and demyelination, whereas WT littermate controls had extensive leukocyte infiltration and demyelination (**Fig. 7F**). T-CTR1-KO mice also had significantly lower levels of CD4^+^ and CD8^+^ T cells, monocytes, macrophages and neutrophils infiltrating their spinal cords (**Fig. 7G**). By contrast, CD4^+^ T cell numbers in the spleen were comparable between WT and CTR1-deficient mice (**Fig. 7H**), suggesting an effect dependent on autoantigen-mediated T cell activation in the CNS. CTR1-deficient CD4^+^ T cells infiltrating the spinal cord had significantly reduced expression of RORγt compared to T-WT cells (**Fig. 7I**). They also showed a significantly impaired production of inflammatory cytokines IL-17A and IFNγ (**Fig. 7J**), as well as GM-CSF (**Fig. S8C**) and IL-2 (**Fig. S8D**). CTR1 was recently reported to be important for the maintenance and anti-inflammatory function of T_reg_ cells^23^. We did not, however, observe significant differences in T_reg_ frequencies in the spinal cords of WT and CTR1-deficient mice with EAE (**Fig. S8E**) or in IL-10 production by CD4^+^ T cells (**Fig. S8F**) suggesting that while CTR1 is essential for the pathogenicity of pT_H_17 cells it appears to be dispensable for T_reg_ cells in the context of EAE. Because untreated T-CTR1-KO mice had a significant decrease in total CD4^+^ T cell numbers in their spleen (although not their LNs) (**Fig. S1D**), we confirmed that their protection from EAE was not due to a decreased global pool of CD4^+^ T cells. To this end, we deleted *Slc31a1* in WT CD4^+^ T cells from 2D2 mice by shRNA (**Fig. S8G-H**) or CRISPR/Cas9 editing (**Fig. S8I-J**), followed by polarization into pT_H_17 cells, adoptive transfer into *Rag1^-/-^*host mice and MOG immunization to induce EAE. Deletion of *Slc31a1* by either shRNA or sgRNA protected host mice from EAE as evidenced by significantly lower EAE disease incidence and severity (**Fig. S8G, I**). Protection from EAE was associated with a reduced infiltration of CD4^+^ T cells (**Fig. S8H, J**), macrophages and neutrophils (**Fig. S8H**) into the spinal cords of host mice, while CD4^+^ T cell numbers in the spleen remained unchanged (**Fig. S8H, J**). Taken together, our data demonstrate that CTR1 is critical for the encephalitogenic function of pT_H_17 cells and EAE development.

## DISCUSSION

### Here we show that CTR1 and copper uptake play an essential role in pT_H_17 cell differentiation and function

We identified CTR1 as a critical regulator of myelin-specific, pT_H_17 cells in the EAE model of MS. Deletion of CTR1 reduces intracellular copper levels in CD4^+^ T cells, confirming its central role in copper import. CTR1 deficiency did not cause major defects in thymic T cell development or in CD4^+^ T cell viability, proliferation, or activation *in vitro*. The strongest phenotype of CTR1 deletion in CD4^+^ T cells was a selective defect in the differentiation and function of pT_H_17 cells associated with reduced expression of RORγt and production of IL-17A and GM-CSF. While homT_H_17 cells also showed reduced cytokine production, their differentiation appeared intact. Importantly, the differentiation and function of T_H_1, T_H_2, and iT_reg_ cells *in vitro* were largely preserved. Transcriptomic analysis reinforced this point. Although we observed differential gene expression in CTR1-deficient T_H_1 and pT_H_17 cells, only core T_H_17-associated genes such as *Il17a/f* and *Il23r* were repressed in the absence of CTR1 whereas the T_H_1 program, including IFN-γ expression, remained comparatively intact. Of note, CTR1-deficient encephalitogenic CD4^+^ T cells isolated from the CNS of mice after EAE induction had reduced IFN-γ levels. A similar apparently paradoxical situation has been described in CD4^+^ T cells with complex I deficiency which can be polarized into Th1 cells and initially produce IFN-γ when metabolic conditions are permissive but fail to sustain continued IFN-γ production once they encounter the metabolically restrictive CNS environment during EAE^52^. The predominant effect of CTR1 deletion on pT_H_17 cells is consistent with the fact that CTR1 expression is highest in pT_H_17 cells relative to other CD4^+^ T cell subsets. The important role of copper in pT_H_17 cells was confirmed using pharmacologic manipulation of copper levels. Copper chelation in WT CD4^+^ T cells phenocopied the pT_H_17 defect observed in the absence of CTR1, whereas copper ionophores rescued the pT_H_17 defect in CTR1-deficient cells.

These results support a model in which CTR1 is not a general checkpoint for T cell function as their activation, viability, and proliferation were mostly preserved. Instead CTR1 acts as a lineage-selective metabolic gatekeeper whose main role is to supply copper needed for pT_H_17 differentiation. This distinction is important because it suggests that copper availability is essential under inflammatory pT_H_17-polarizing conditions driven by IL-1β, IL-6, and IL-23. The high CTR1 expression in pT_H_17 cells implies that these cells either require more copper than other subsets or are more sensitive to reductions in copper-dependent processes. These findings are intriguing in light of a recent study showing that CTR1 is essential for T_reg_ cells, with T_reg_-specific deletion of CTR1 resulting in pathological autoinflammation in mice^23^. Like pT_H_17 cells, T_reg_ cells rely on mitochondrial respiration to exert their immunoregulatory functions^53^. In our hands, T cell-specific deletion of CTR1 did not impair iT_reg_ differentiation in vitro or cause an autoimmune phenotype in vivo. The lack of overt autoimmunity is likely explained by the defective function of pT_H_17 cells. Another study demonstrated that haploinsufficiency for CTR1 in CD4^+^ T cells results in increased production of IL-2, IFN-γ, and TNF due to reduced copper levels, enhanced histidine phosphorylation of the potassium channel K_Ca_3.1, and thus increased calcium influx^54^. We did not observe augmented production of these cytokines by CTR1-deficient T_H_ cell subsets *in vitro* or *in vivo*. A possible explanation for this difference could be the gradual effects of strong copper depletion in our study, compared with the more moderate reduction in CTR1^+/-^ CD4^+^ T cells reported earlier^54^.

Our finding that CTR1 functions as a selective regulator of pT_H_17 cell fate expands the concept that trace metals are not merely supportive nutrients but active determinants of immune cell identity. Our findings also have therapeutic relevance because inhibiting CTR1 function, or copper import more generally, in CD4^+^ T cells suppresses the differentiation and encephalitogenic effector functions of pT_H_17 cells while sparing other T_H_ cell subsets and broader CD4^+^ T cell functions. Targeting copper uptake by CTR1 could therefore represent a mechanistically grounded strategy for reducing pT_H_17 cell function without the broad immunosuppression associated with many current therapies.

### Mechanistically, we propose a model that links copper uptake by CTR1 to mitochondrial metabolism and the epigenetic regulation of pT_H_17 differentiation and function

CTR1-deficient cells have reduced intracellular copper levels, which is associated with decreased mitochondrial ROS production and respiration, consistent with the known role of copper in COX (complex IV) function^16^. Our metabolomics analyses further showed an accumulation of dihydroorotate and glycerol-3-phosphate, accompanied by a reduced NAD^+^/NADH ratio and compensatory increases in glycolysis, suggesting that impaired oxidative metabolism is offset by glycolytic rewiring. Importantly, the defect in mitochondrial metabolism is functionally linked to pT_H_17 differentiation. Inhibition of mitochondrial ATP synthase with oligomycin reduced pT_H_17 differentiation and cytokine production in WT cells, while T_H_1 cells were comparatively resistant. These findings corroborate the notion that pT_H_17 cells exhibit a particular dependence on mitochondrial respiration, which contributes to the epigenetic regulation gene expression, consistent with previous observations^5,25–29^.

While mitochondrial respiration was shown to promote pT_H_17 differentiation and function, the mechanisms underlying this regulation are not fully understood. Here, we provide a copper-dependent link between mitochondrial metabolism and the T_H_17 transcriptional program. We propose a model in which copper depletion blocks mitochondrial respiration, resulting in the accumulation of TCA intermediates that are potent inhibitors of TET enzymes. CTR1-deficient pT_H_17 cells accumulated 2-hydroxyglutarate, succinate, and fumarate, which is intriguing because all three metabolites inhibit TET DNA demethylases and histone demethylases. Consistent with impaired demethylation, CTR1-deficient cells displayed increased global DNA methylation and hypermethylation of key T_H_17-associated genes. These methylation changes correlated with lower chromatin accessibility and reduced expression of T_H_17 signature genes. Direct functional perturbation of DNA methylation further supported these findings: treatment of WT CD4^+^ T cells with 2-HG or a TET inhibitor impaired pT_H_17 differentiation and function, whereas DNMT inhibitors rescued the expression of RORγt, IL-17A, and GM-CSF in CTR1-deficient T cells. In line with our findings, DNA methylation was previously reported to regulate *Il17a* transcription in T_H_17 cells in vitro^48,49^, and in T cells from patients with systemic lupus erythematosus^55^. Moreover, histone methylation was reported to regulate pT_H_17 proliferation, cytokine production, and pathogenicity in EAE^56^. While not tested directly in our study, we speculate that the metabolic rewiring of CTR1-deficient T cells may also perturb histone methylation and thus pT_H_17 differentiation. Intriguingly, increased H3K27me3 was recently described in CTR1-deficient T_reg_ cells^23^.

Our study also provides evidence of widespread changes in chromatin accessibility in CTR1-deficient pT_H_17 cells. Motif enrichment and TF footprinting analyses of ATAC-seq data predicted reduced AP-1 family transcription factor binding, especially BATF, JunB, and Fosl2, at T_H_17-associated gene loci such as *Il17a/f* and *Il21*. While the precise roles of these TFs in copper-dependent regulation of pT_H_17 differentiation remain to be elucidated, our data suggest that epigenetic dysregulation in CTR1-deficient T cells alters the recruitment of TFs that play pioneering roles in establishing the T_H_17 program^43,44^. In addition to the depletion of binding motifs for TFs that promote T_H_17 differentiation, our analysis also showed enrichment of motifs for TFs that repress T_H_17 differentiation such as Ets1^47^. Ets1 is involved in T_H_ cell specification by interacting with BATF and promoting the recruitment of CCCTC-binding factor (CTCF), a zinc finger protein regulating 3D chromatin structure^43^. Although CTCF was shown to modulate T_H_17 cell differentiation^43^, a potential involvement of CTR1 in this pathway remains speculative and beyond the scope of this study.

Our data link copper and mitochondrial metabolism to the epigenetic regulation of pT_H_17 differentiation and function. We propose a model in which limited copper uptake in the absence of CTR1 results in impaired mitochondrial respiration and the buildup of metabolites that inhibit demethylating enzymes. The resulting DNA hypermethylation creates an epigenetic bottleneck that prevents the proper chromatin opening and the recruitment of lineage-defining TFs required for pT_H_17 differentiation. This model is not simply an energy-deficit model linked to impaired ATP production as CTR1-deficient CD4^+^ T cells remain viable, proliferate, and can be activated. Instead, we propose that CTR1 and copper sustain a metabolic state permissive for epigenetic remodeling, and that pT_H_17 cell fates are governed by metabolic intermediates that epigenetically regulate their differentiation.

### The critical role of CTR1 and copper in pT_H_17 cells has significant translational implications

If copper availability shapes pathogenic gene expression programs by controlling DNA methylation and chromatin accessibility, then therapeutic strategies targeting copper uptake, mitochondrial metabolism, or downstream epigenetic programs might be used to modulate pT_H_17 cell function in autoimmunity and inflammation. In patients with relapsing-remitting MS, circulating effector T cells expressed higher *SLC31A1* levels than those from healthy controls. In the EAE model of MS, *Slc31a1* expression was increased in CNS-infiltrating CD4^+^ T cells relative to splenic CD4^+^ T cells potentially indicating an increased demand for copper-dependent mitochondrial function in encephalitogenic T cells. Functionally, T cell-specific deletion of CTR1 strongly protected mice from EAE, as reflected by attenuated disease incidence and severity. Moreover, CNS inflammation and demyelination were markedly reduced in the absence of CTR1. In particular, autoreactive CD4^+^ T cells were almost absent from the CNS, and those present showed strongly reduced production of IL-17A, IFN-γ, and GM-CSF. Importantly, the numbers of CTR1-deficient CD4^+^ T cells in the spleens of mice with EAE were not reduced. Our study identifies CTR1 as a potentially actionable pathway for the treatment of autoimmune neuroinflammation in MS by exploiting the copper-dependent metabolic liability of pT_H_17 cells. While these findings do not yet establish clinical efficacy or safety in humans, they provide a rationale for exploring copper-targeted approaches in MS and possibly other T_H_17-driven autoimmune diseases. Additional support for such a strategy comes from reports that copper levels are increased in patients with autoimmune diseases^17–20^ and that the copper chelator TTM has protective effects in the murine EAE model^21^ and in a rat model of RA^57^. Moreover, the copper chelator LCC-12 is a potent inhibitor of inflammation in mice^22^, suggesting that copper depletion could be a promising avenue for the treatment of autoimmune diseases. When considering copper-targeting treatments in CNS inflammation, however, other effects of copper must also be considered. Astrocytes were shown to release copper into the extracellular milieu in response to inflammation and to contribute to demyelination and oligodendrocyte loss^58^. Moreover, the copper-chelating toxin cuprizone is widely used to model demyelination and oligodendrocyte loss in MS^59^. Copper is also critical for neuronal metabolism and protection against oxidative stress, and copper deficiency results in neurological impairments^60^.

Collectively, our findings position CTR1-dependent copper uptake as a key metabolic and epigenetic driver of pT_H_17 responses and a potential therapeutic target in MS. Going forward, carefully designed studies must determine whether copper-modulating strategies can safely suppress autoimmune neuroinflammation without disrupting the essential roles of copper in the CNS.

## METHODS

### Mice

All mouse strains were purchased from The Jackson Laboratory (JAX): C57B6/J (JAX 000664), CD45.1 (JAX 002014), *Rag1*^-/-^ (JAX 002216), CD4Cre (JAX 022071), 2D2 (JAX 006912), Cas9^GFP^ (JAX 026179), *Slc31a1^F^*^/F^ (*Slc31a1^tm2Djt^*/J; JAX 025651). Double transgenic 2D2-Cas9^GFP^ mice were generated by crossing 2D2 and Cas9^GFP^ mice. To obtain T-cell specific deletion of *Slc31a1* (T-CTR1-KO mice), *Slc31a1^F^*^/F^ mice were crossed with CD4Cre mice. All mice were maintained under pathogen-free conditions at 22-25°C, 50-60% humidity, with a 12-hour dark/light cycle, and food and water *ad libitum*. Age and sex-matched female and male mice of 6 to 20 weeks of age were used for experiments. All experiments were conducted in accordance with protocols approved by the Institutional Animal Care and Use Committee of New York University Grossman School of Medicine.

### T cell culture

Primary mouse CD4^+^ T cells were isolated from the spleens of mice using the EasySep Mouse CD4^+^ T cell Isolation Kit (StemCell Technologies; 19852), following the manufacturer’s instructions. Naïve CD4^+^ T cells were stimulated for 2 days with 1 µg/ml plate-bound anti-mouse CD3ε and CD28 antibodies in tissue culture-treated plates pre-coated with Rabbit anti-Syrian Hamster IgG (25 µg/ml in PBS) (the complete list of antibodies used in this study can be found in **Table S1**). Cells were cultured in RPMI1640 tissue culture medium (Corning; 10-040-CV) supplemented with 10% FBS (Sigma-Aldrich; F0926-500ML (17A412)), 2mM L-Glutamine (ThermoFisher; 25030081), 50 U/ml penicillin/streptomycin (ThermoFisher; 15140163), 50µM β-mercaptoethanol (MP Biomedicals; ICN19024280). After 2 days of stimulation, T cells were detached and transferred to a new plate with fresh complete RPMI medium with 20 U/ml rhIL-2 (PeproTech; 200-02), except for T_H_17 culture conditions. For differentiation into pT_H_17, cells were cultured in the presence of 5 µg/ml anti-IFN-γ, 5 µg/ml anti-IL-4, 20 ng/ml IL-6 (R&D Systems; 406-ML), 20 ng/ml IL-1β (PeproTech; 211-11B) and 20 ng/ml IL-23 (R&D Systems; 11269-ML). For differentiation into homT_H_17, cells were cultured in the presence of 5 µg/ml anti-IFN-γ, 5 µg/ml anti-IL-4, 20 ng/ml IL-6 and 0.5 ng/ml hTGFβ (PeproTech; 100-21). For differentiation into T_H_1, cells were cultured in the presence of 5 µg/ml anti-IL-4, 20 ng/ml IL-12 (PeproTech; 210-12). For differentiation into T_H_2, cells were cultured in the presence of 5 µg/ml anti-IFN-γ and 50 ng/ml IL-4 (PeproTech; 214-14). For differentiation into iT_reg_, cells were cultured in the presence of 5 µg/ml anti-IFN-γ, 5 µg/ml anti-IL-4 and 5 ng/ml hTGFβ.

### Transfection and retroviral transduction of shRNA and sgRNA

The shRNA EAE screen for ICTs in encephalitogenic T cells was described previously^7,8^. shRNA target sequences against *Slc31a1* were extracted from the shRNA library, shRNA targeting Renilla luciferase was used as control. shRNAs were cloned into the pLMPd retroviral expression vector with either Thy1.1 or Ametrine reporter^61^. sgRNA target sequences were designed using CRISPick sgRNA Designer (Broad Institute, Cambridge, MA)^62^, an sgRNA targeting human VEGFR was used as control. sgRNAs were cloned intro the pMRI expression vectors with Ametrine reporter^63^. All shRNA and sgRNA sequences are provided in **Table S2**. Platinum-E (Plat-E) retroviral packaging cells were cultured in DMEM medium (Corning; 10-013-CV) supplemented with 10% FBS, 1 μg/ml of puromycin dihydrochloride (InvivoGen; A11138-03), 10 μg/ml of blasticidin (InvivoGen; Ant-bl-1) and 50 U/ml of penicillin/streptomycin. For retrovirus production, Plat-E cells were grown to 80% confluency and transfected using GenJet lipofection reagent (SignaGen; SL100489) with 1.5 µg of retroviral expression plasmids and 0.6 µg ecotropic packaging vector pCL-10A1; 0.6 µg of shPasha plasmid was added for shRNA vectors (per well in 6 well plate). Media was changed 16h after transfection, and retroviral supernatant was collected 48 and 60 hours after transfection. For retroviral transduction of mouse CD4^+^ T cells, cells were stimulated for 20-24 hours as described above, then media was replaced with retroviral supernatant and 8 μg/ml of polybrene (hexadimethrine bromide; Sigma-Aldrich; 107689), followed by centrifugation of cells at 1450 x *g* for 90 min at 32°C for spin-infection.

### Flow cytometry and cell sorting

Cells were washed in PBS containing 3% FBS (FACS buffer). Cells were first incubated with anti-CD16/32 (BioXcell; BP0307) for 10 minutes. Staining of surface molecules with fluorescent labeled antibodies was performed in FACS buffer for 15 minutes. The complete list of antibodies and dilutions used in this study can be found in **Table S1**. Dead cells were labeled using either 0.5 µM DAPI (Cayman Chemical; 14285) or LIVE/DEADTM Fixable Blue Dead Cell Stain Kit (Invitrogen; L34962 A) following the manufacturer’s instructions. Surface CTR1 protein detected on live cells with a primary antibody targeting the extracellular N-terminus portion of CTR1, followed by a fluorescently labeled secondary antibody. To measure intracellular copper levels, cells were loaded with CD649.2 Cu(II) dye as previously described^24^. To measure cell proliferation, cells were loaded with CellTrace Violet (CTV) (ThermoFisher Scientific; C34571) according to the manufacturer’s instructions, and dye dilution was monitored daily. To measure cell viability, cells were stained Alexa Fluor 647 Annexin V (BioLegend; 640912) at 1:33 dilution for 10 minutes in Annexin V binding buffer (BioLegend; 422201). To measure mitochondrial ROS production and mitochondrial content, cells were stained with 1 µM MitoSOX Red (Invitrogen; M36008), 100 nM MitoView Green (Biotium; 70054), MitoTracker Deep Reed (Invitrogen; M46753) according to the manufacturer’s instructions. To measure total ROS levels, cells were incubated with 5 µM H_2_DCFDA (ThermoFisher; D399) for 30 minutes at 37C. To analyze cytokine production, CD4^+^ T cells were stimulated with 20 nM phorbol myristate acetate (PMA; Calbiochem; 524400) and 1 µM ionomycin (Invitrogen; I-24222) or 1:500 stimulation cocktail (containing 0.0405 mM PMA and 0.67 mM ionomycin calcium salt; eBioscience; 00-4970-93eBioscience; 00-4970-93), in the presence of 3 µg/ml brefeldin A (eBioscience; 00-4506-51) in complete media for 4 hours. Cells were then collected, washed in FACS buffer and fixed using IC fixation buffer (Invitrogen; 00-8222-49), following the manufacturer’s instructions. Cells were then incubated in permeabilization buffer (Invitrogen; 00-8333-56), stained with antibodies for 1 hour at room temperature and washed. Transcription factor staining was performed using 1 part of TF fixation buffer concentrate (eBioscience; 00-5123-43) diluted into 3 parts of diluent (eBioscience; 00-5223-56), following the manufacturer’s instructions. Cells were then incubated with permeabilization buffer, stained for 1 hour at room temperature and washed. Samples were acquired on an LSR Fortessa flow cytometer (BD Biosciences) and analyzed using FlowJo software (BD Biosciences; version 10.10.1). Total cell counts were obtained with Precision Count Beads (BioLegend; 424902) following the manufacturer’s instructions. Enrichment of CD4^+^ T cells by flow cytometric cell sorting was performed with a FACSAria II (BD Biosciences) using a 70 μm nozzle.

### Experimental Autoimmune Encephalomyelitis (EAE)

To induce active EAE T-WT or T-CTR1-KO mice were immunized with 0.2 ml MOG_35-55_ emulsion in complete Freund’s adjuvant (CFA) (Hooke Laboratories, EK-2110), followed by intraperitoneal injection of 150 ng/ml Pertussis toxin (Hooke Laboratories, EK-2110, lot #1015) at 2 and 26 hours after MOG immunization. To induce passive EAE, CD4^+^ T cells were isolated from the spleens and lymph nodes (LNs) of 2D2 mice, activated and differentiated into pT_H_17 cells as described above. T cells were transduced with shRNA at 24 hours after activation. After 24 hours, cells were detached and cultured in fresh complete media for an additional 48 hours. Transduced cells were enriched by flow cytometric cell sorting and 200,000 cells diluted in 100 µl PBS were adoptively transferred into *Rag1*^-/-^host mice by retro-orbital injection. For *in vivo* T cell competition assay, CD4 T cells from 2D2 mice transduced with either shControl plasmid (containing an Ametrine reporter) or sh*Slc31a1* plasmid (containing an Ametrine reporter) were mixed at a 1:1 ratio with 2D2 T cells transduced with shControl plasmid (containing a Thy1.1 reporter) and injected into *Rag1^-/-^* mice. After 2 days, host mice were immunized with 200 μg of MOG_35-55_ peptide (Anaspec; AS-60130-1) emulsified in CFA (Difco; DF0638-60-7) with H37Ra (Difco; 231141). The severity of EAE was scored in a blinded manner according to the following disease scoring system as described^64^: 0 = no disease; 0.5 = partially limp tail; 1 = paralyzed tail; 2 = hindlimb weakness; 2.5 = partial hindlimb paralysis; 3 = hindlimb paralysis; 3.5 = hindlimb paralysis and hunched back; 4 = hindlimb and forelimb paralysis; 5 = moribundity and death. All animals were supported with DietGel 76A (ClearH2O; 72-07-5022). Mice were euthanized when they lost more than 20% of their original body weight or showed hindlimb and forelimb paralysis. At the end of the experiment, cells were isolated from the spinal cord and spleen of mice and analyzed by flow cytometry. Spinal cords were minced and digested with 1 mg/ml collagenase D (Sigma-Aldrich; 11088882001) with gentle agitation for 45 min at 37 °C. Spinal cord homogenates were filtered through a 70 μm strainer, and immune cells were enriched using a 38% Percoll (Sigma-Aldrich; P4937) gradient and centrifugation at 900 x *g* for 20 minutes at room temperature.

### Inductively Coupled Plasma Mass Spectrometry (ICP-MS)

CD4^+^ T cells were isolated from either T-WT or T-CTR1-KO mice and polarized under pT_H_17 conditions for 4 days as described above. 2.5 x 10^6^ cells were collected, washed and lysed in nitric acid (VWR Chemicals BDH Aristar Ultra, 87003-228) at room temperature on a rotator in 1.5 mL tubes (Sarstedt). Digested samples were diluted in 2% nitric acid with 20 p.p.b. Gallium solution (inorganic Ventures, AAGA1-125ML) as internal standards and analyzed with iCAP-Qc ICP-MS (Thermal Fisher).

### RNA sequencing

CD4^+^ T cells were isolated from either T-WT or T-CTR1-KO mice, activated and polarized in either pT_H_17 or T_H_1 conditions for 3 days as described above. Total RNA was extracted from 1 x 10^6^ T cells using the RNeasy Mini RNA Isolation Kit (Qiagen; 74104). RNA quality and quantity were analyzed using a Nanochip cartridge and the Agilent 2100 Bioanalyzer system. RNAseq libraries were prepared using the Illumina TruSeq stranded total RNA kit (Illumina; RS-122-2202), starting from 100 ng of DNAse I (Qiagen)–treated total RNA following the manufacturer’s protocol with 15 PCR cycles. The amplified libraries were purified using AMPure beads (Beckman Coulter; A63882), quantified using a Qubit 2.0 fluorometer (Life Technologies), and visualized using the high-sensitivity DNA ScreenTape on the Agilent TapeStation 2200 system. The libraries were pooled equimolarly and analyzed by paired-end sequencing using a P2 100 cycle flow cell and the Illumina NextSeq 2000 system. Fastq files were created from Illumina output via bcl2fastq2 Conversion software (v2.20). Illumina adaptors were trimmed using Trimmomatic (v0.36)^65^. Assembly was performed with GTF reference assistance with the trimmed reads using STAR (v2.7.7), with mm10 as the reference genome. The resulting alignment maps were converted into a gene count matrix using the Feature count function of Bed tools (v2.10). Differential gene expression was analyzed using DESeq2 (v1.34.0)^66^ in an R statistical programming environment. Genes with an expression of less than 10 counts in half of the samples were excluded from further analysis. Differentially expressed genes (DEGs) were considered significant when adjusted *P* values were ≤ 0.01. Log-fold shrinkage was calculated using the native APEGM function of DESeq2^67^. Functional pathway analyses of DEGs were performed using the GSEA function of ClusterProfiler 4.0^68^. We further combined the canonical databases of the Molecular Signatures Database (MSigDB), along with all three of the gene ontologies and the Hallmark GSEA pathways datasets^69^, to create a robust pathway database. Genes found to be significantly dysregulated were also used to conduct a pathway analysis to confirm GSEA findings. The analysis of upstream regulators of DEGs was conducted using QIAGEN Ingenuity Pathway Analysis (Qiagen)^70^. Pathways were considered significant if they had a Bonferroni adjusted *P* value of ≤ 0.01. GSEA plots were generated using the gseaplot2 function of clusterProfiler. Enrichment network maps were calculated using the pairwise function in clusterProfiler, and the resulting similarity matrix was visualized using igraph^71^. Edges between pathways were removed if they did not meet a threshold of 30% shared genes.

### Chemicals

All chemicals used in this study were prepared and stored following manufacturer’s recommendations. Chemicals used for chronic treatment during T cell activation and differentiation were titrated to find the threshold that does not compromise viability and activation of CD4^+^ T cells in vitro. All titration data can be found in **Fig. S9** and **S10**. The following chemicals were used for this study, following conditions as indicated in corresponding figure legends: Tetrathiomolybdate (TTM; Sigma-Aldrich; 323446), LCC-12 (Cayman Chemical; 38929), CuGTSM (Chang lab), Elesclomol (Cayman Chemical; 10011192), Oligomycin A (Cayman Chemical; 11342), LCS-1 (Cayman Chemical; 35231), Tiron (Abcam; ab146234), Chloroquine diphosphate salt (Sigma-Aldrich; C6628), Dimethyl succinate (Sigma-Aldrich; 8201500250), Monomethyl fumarate (Cayman Chemical; 27813), (2R)-Octyl-α-hydroxyglutarate (2-HG; Cayman Chemical; 16366), 5-azacytidine (Cayman Chemical; 11164), Decitabine (Cayman Chemical; 11166), TETi76 diethyl ester (Sigma-Aldrich; SML3121), SAHA (Cayman Chemical; 10009929), Trichostatin A (TSA; Cayman Chemical; 89730), Anacardic Acid (Cayman Chemical; 13144), CPTH2 (Cayman Chemical; 12086), A-485 (Cayman Chemical; 24119).

### Extracellular flux analysis

CD4^+^ T cells were analyzed after 1 or 3 days of activation and culture in either T_H_1 or pT_H_17 conditions, as previously described. On the day of the analysis, 2.5 x 10^5^ cells were seeded per well in 24 well plates pre-coated with 0.01% poly-L-lysine (Sigma-Aldrich; P8920) in triplicate in XF DMEM assay medium (Agilent; 103680-100). Cells were analyzed with the XF Mito Stress Test Kit (Agilent; 103015-100), the XFe24 FluxPak (Agilent; 102340-100) and the XFe24 Analyzer (Agilent) using 1 µM oligomycin, 1.5 µM FCCP and 0.5 µM Rotenone/Antimycin A, following manufacturer’s instructions.

### Liquid Chromatography-Mass Spectrometry (LC-MS) metabolomics

CD4^+^ T cells were isolated form T-WT or T-CTR1-KO mice, activated and differentiated under pT_H_17 polarizing conditions for 3 days as described above. Cells were pelleted for 30 sec at 3000 x g, washed in ice cold PBS, pelleted at 3000 x g for 30 sec, and resuspended in ice cold lysis buffer containing 80% methanol (Fisher Scientific; 67-56-1) and 500 nM metabolomics amino acid mix (Cambridge Isotope Laboratories; MSK-A2-1.2) at 5 x 10^6^ cells/ml. Samples were vortexed in 2 ml screw cap vials containing ∼100 µL of disruption beads (Research Products International), then subsequently spun at 21,000 g for 3 min at 4 °C. A set volume of each (450 µL) was transferred to a 1.5 ml tube and dried down by speedvac (Thermo Fisher). Samples were reconstituted in 50 µl of Optima LC/MS grade water (Fisher Scientific). Samples were sonicated for 2 mins, then spun at 21,000 g for 3 min at 4°C. 20 µl were transferred to LC vials containing glass inserts for analysis. The remaining sample was placed in -80 °C for long term storage. Samples were subjected to an LCMS analysis to detect and quantify known peaks, as previously described^72^. The LC column was a MilliporeTM ZIC-pHILIC (2.1 x150 mm, 5 μm) coupled to a Dionex Ultimate 3000TM system and the column oven temperature was set to 25 °C for the gradient elution. A flow rate of 100 μl/min was used with the following buffers: A) 10 mM ammonium carbonate in water, pH 9.0, and B) neat acetonitrile. The gradient profile was as follows; 80-20%B (0-30 min), 20-80%B (30-31 min), 80-80%B (31-42 min). Injection volume was set to 2 μl for all analyses (42 min total run time per injection). MS analyses were carried out by coupling the LC system to a Thermo Q Exactive HFTM mass spectrometer operating in heated electrospray ionization mode (HESI). Method duration was 30 min with a polarity switching data-dependent Top 5 method for both positive and negative modes. Spray voltage for both positive and negative modes was 3.5 kV and capillary temperature was set to 320 °C with a sheath gas rate of 35, aux gas of 10, and max spray current of 100 μA. The full MS scan for both polarities utilized 120,000 resolution with an AGC target of 3e6 and a maximum IT of 100 ms, and the scan range was from 67-1000 m/z. Tandem MS spectra for both positive and negative mode used a resolution of 15,000, AGC target of 1e5, maximum IT of 50 ms, isolation window of 0.4 m/z, isolation offset of 0.1 m/z, fixed first mass of 50 m/z, and 3-way multiplexed normalized collision energies (nCE) of 10, 35, 80. The minimum AGC target was 1e4 with an intensity threshold of 2e5. All data were acquired in profile mode.

### Single cell energetic metabolism by profiling translation inhibition (SCENITH)

CD4^+^ T cells were isolated from T-WT or T-CTR1-KO mice, activated and differentiated under pT_H_17 polarizing conditions for 3 days as described above. To assess the cells’ dependence on glucose, cells were collected and treated with 50 mM 2-deoxyglucose (2-DG; Cayman Chemical; 14325) or vehicle control for 15 minutes, and in the presence of 10 µg/ml puromycin (Invitrogen; A11138-03) for an additional 30 minutes. Cells were washed, fixed, permeabilized and stained for puromycin for flow cytometry analysis as described above. Glucose dependence was defined as the difference in puromycin mean fluorescence intensity (MFI) between vehicle control and 2-DG-treated cells.

### Real-time quantitative PCR

RNA was isolated from CD4^+^ T cells using TRIzol reagent (Invitrogen; 15596026) and quantified using a Nanodrop 8000 spectrophotometer (Thermo Scientific). cDNA was synthesized using the iScript™ cDNA synthesis kit (Bio-Rad; 1708891). Quantitative real-time PCR was performed using the Maxima SYBR Green qPCR Master Mix (ThermoScientific; K0223). Transcripts levels were normalized to the expression of *Rpl32* housekeeping gene using the 2^−ΔCT^ method. Data was acquired using QuantStudio Design & Software Analysis using a QuantStudio 3 PCR instrument (Applied Biosystem). A complete list of primers is provided in **Table S2.**

### Immunoblot analysis

CD4^+^ T cells were washed with ice-cold PBS, pelleted and lysed in RIPA buffer (Sigma-Aldrich; R0278) supplemented with protease inhibitor cocktail (Sigma-Aldrich; P8340) and phosphatase inhibitor cocktail (Cell signaling; 5870S). Cell lysates were precleared by centrifugation at 16,000 x *g* for 10 min at 4°C, and cleared lysates were resuspended in Laemmli buffer (Biorad; 1610747). Samples were resolved on 4-20% SDS-PAGE gels (Biorad; 4561096) and transferred to PVDF membranes (BioRad; 162-0184). Membranes were blocked with 5% nonfat milk (ThermoFisher; sc-2325) for 1 hour at room temperature and incubated with primary antibodies overnight at 4°C with gentle rocking. Membranes were washed three times for 10 minutes with Tris-buffered saline containing 0.1% Tween 20 (TBST; American Bioanalytical; AB02038) followed by incubation with corresponding HRP-conjugated secondary antibodies for 1 hour at room temperature. The reaction was visualized using chemiluminescent ECL reagent Clarity Max (BioRad; 1705062) and detected using an Amersham Imager 680. All antibodies and dilutions can be found in **Table S1.**

### NAD^+^, NADH measurements

CD4^+^ T cells were isolated from either T-WT or T-CTR1-KO mice and activated under pT_H_17 polarizing conditions for 3 days as described above. For the assay, 8 x 10^5^ cells were collected and analyzed using the NAD/NADH-Glo assay (Promega; G9071), following the manufacturer’s instructions and a FlexStation 3 plate reader (Molecular Devices).

### SOD activity measurements

CD4^+^ T cells were isolated from either T-WT or T-CTR1-KO mice and activated for 3 days as described above. For the assay, 1 x 10^6^ cells were collected and processed with the SOD activity assay (Sigma-Aldrich; CS0009), following the manufacturer’s instructions, and measured with a FlexStation 3 plate reader.

### Global 5mC measurements

CD4^+^ T cells were isolated from either T-WT or T-CTR1-KO mice, activated under pT_H_17 polarizing conditions for 3 days as described above. To measure global 5mC levels, 5 x 105 cells were collected and processed with the MethylFlash Global DNA methylation (5mC) ELISA Easy kit (Epigentek; P103096), following the manufacturer’s instructions, and analyzed using a FlexStation 3 plate reader.

### DNA methylation sequencing

CD4^+^ T cells were isolated from either T-WT or T-CTR1-KO mice, activated under pT_H_17 polarizing conditions for 3 days as described above. High molecular weight DNA was extracted from 1 x 10^6^ cells using the Nanobind CBB kit (PacBio; 102-207-600), following the manufacturer’s instructions.

For Nanopore long read sequencing, 3 µg of genomic DNA from each sample was sheared with a Covaris G-tube to an average base pair size of 20-30kb in a tabletop Eppendorf centrifuge. The sheared DNA was input into the Oxford Nanopore Ligation Sequencing Kit v14 (SQK-LSK114) to create a nanopore library according to manufacturer’s protocol with extended incubation times of 1 hour for each step. The final library was eluted with buffer overnight at room temperature to allow for larger pieces of DNA to elute off the ampure beads. The library samples were prepared without PCR to preserve the methylation across the genome. The final library was verified on Qubit using the High Sensitivity DNA kit to assess the loading concentration. Each sample was sequenced on 1 Promethion flow cell R10 (FLO-PRO114M) with methylation calls enabled for both 5mC & 5hmC (5-hydroxymethylcytosine). Raw sequencing reads were basecalled using the high-accuracy (HAC) Dorado basecalling function (v1.2.0)^73^ and aligned to mm10 reference assembly. Aligned reads were sorted, indexed, prior to downstream methylation analysis using samtools (v1.9). Cytosine modification calls were processed using modkit (v0.4.4)^74^. Modification calls were exported in bedMethyl format for downstream differential methylation and genomic annotation analyses, using the dmr pair function. The regions of interest were defined as listed below.

Alternatively, DNA libraries were prepared with the Biomodal Duet evoC library prep kit starting from 150 ng of gDNA for each sample (Biomodal 8rxn, #6205/4103, version 4 protocol 2024). Libraries were quantified with the Kapa-Roche Complete Universal qPCR kit to confirm nanomolar concentration. Libraries were normalized and pooled equimolar into a final pool for sequencing. The pool was sequenced on the Illumina Novaseq X-Plus system using 1 lane of the 25B 300 cycle flow cell. Sequencing metrics were run as paired end 150 bases. Raw sequencing data were processed using the biomodal duet software pipeline to resolve canonical nucleotide sequence and cytosine modification states. Reads were aligned to the mm10 mouse reference genome, and 5mC as well 5hmC levels were quantified at single-base resolution. Downstream methylation analyses were performed using the biomodal modality XPLR software. Differentially methylated regions (DMRs) were identified using the modality DMR core analysis workflow (v1.1). Genome-wide 5mC counts were aggregated across predefined genomic regions, and the numbers of methylated and total cytosines were summed for each sample within each region. Regions assessed for differential methylation included 1kb windows, gene bodies spanning the 5′ untranslated region (UTR) through the 3′ UTR, promoter regions encompassing the transcription start site and 3 kb upstream and downstream of the transcription start site, and gene-associate enhancer regions defined using ENCODE Project annotations v4^75^.

### Assay for Transposase-Accessible Chromatin with sequencing (ATACseq)

CD4^+^ T cells were isolated from T-WT or T-CTR1-KO mice and activated under pT_H_17 polarizing conditions for 3 days. ATAC-seq was performed according to the manufacturer’s protocol (Active Motif; 53150). Briefly, 1 x 10^5^ T cells were washed in ice-cold PBS and lysed in ATAC-seq lysis buffer. Cell nuclei were pelleted by centrifugation at 500 x *g* for 10 min at 4°C. After removal of the supernatant, nuclei were resuspended in Tagmentation Master Mix and incubated for 30 min at 37°C. DNA was isolated using a DNA purification buffer and columns. Purified DNA was subjected to PCR amplification with different combinations of i5 and i7 indexed primers.

Amplified DNA was cleaned up using SPRI beads. DNA libraries were pooled equimolarly and analyzed by paired-end sequencing using a SP100 NovaSeq 6000 flow cell and the Illumina NovaSeq 6000 Sequencing System. FASTQ files were generated using the bcl2fastq2 Conversion software (v2.20). Adaptor removal and quality control read trimming was completed using Trimmomatic (v0.36) and the FastQC package was used to confirm sample quality. ATACseq samples were aligned using bowtie2 (v2.3.4.1)^76^. The resulting BAM files were deduplicated using Sambamba (v0.6.8)^77^. Peaks were called using MACS2 (v2.1.1)^78^ and annotated using ChIPseeker (v1.30.3)^79^. Differential accessibility was quantified using the DiffBind package^80^. Tracks were converted into bigwigs using Bedtools (v2.10) for visualization and normalized to reads per kilobase per million.

### Transcription Factor Footprinting Analysis and Motif Enrichment

To identify transcription factor binding motifs associated with differentially accessible chromatin regions, motif enrichment analysis was performed using HOMER (v5.1)^81^. Differentially accessible peaks were analyzed using the findMotifsGenome.pl function against the mm10 reference genome. Regions of interest were compared to GC-content–matched genomic background regions generated by HOMER. Opening and closing regions were separated and both known and de novo motif enrichment analyses were performed using default parameters for each group. Significantly enriched motifs were ranked according to hypergeometric p-values. To infer changes in transcription factor occupancy, foot printing analysis was conducted using TOBIAS^82^. Replicate ATAC-seq BAM files and peak sets were combined and processed using the TOBIAS ATACorrect module to correct for Tn5 transposase insertion bias. Corrected insertion signals were subsequently used to calculate transcription factor footprint scores with the ScoreBigwig module. Differential transcription factor binding activity between experimental groups was assessed using the BINDetect module with position weight matrices obtained from the HOCOMOCO motif database(v11). Changes in footprint depth and flanking accessibility were integrated to estimate differential transcription factor occupancy. Transcription factors exhibiting significant changes in binding scores were identified according to TOBIAS default statistical criteria and visualized using volcano plots and footprint aggregate profiles.

### Histology

Sections of spinal cord were fixed in 4% paraformaldehyde in PBS (Santa Cruz; sc-281692), imbedded in paraffin and cut into 5 μm sections. Slides were stained with hematoxylin and eosin (H&E) or 0.1% Luxol Fast Blue (Acros, Cat. No. 1328-51-4) in 95% ethanol using standard methods. Images were acquired using a Leica AT2 slide scanner and visualized using OMERO (The Open Microscopy Environment).

### Statistical analyses and study design

Mice used for in vivo experiments were assigned to groups randomly. Sample sizes were predetermined to yield statistically significant differences based on the anticipated effects sizes. EAE histology samples were analyzed in blinded. The statistical significance of differences between experimental groups was determined by unpaired Student’s *t* test, paired Student’s *t* test, Welch’s *t* test, Mann-Whitney test, mixed-effects modeling, one-way ANOVA, 2way ANOVA, as indicated in the figure legends. The number of mice per experimental group are indicated in the figure legends. All results are shown as means ± SEM. Normal distribution was tested in experiments with n > 20, using the D’Agostino & Pearson test, Anderson-Darling test, Shapiro-Wilk test and Kolmogorov-Smimov test. All bioinformatics analyses were performed in an R statistical environment (v4.1.1). For ATAC-seq and RNA-seq analyses we used the two-tailed Student’s *t* test adjusted using the Benjamini-Hochberg method. For methylation analysis statistical significance was determined using a Wald test, and P values were corrected for multiple hypothesis testing using the Benjamini–Hochberg method. Differences were considered significant for *P* values < 0.05.

## Acknowledgements

We thank Dr. P. Schwarzberg (NIH) for providing pMRI-Amt and pMRI-GFP plasmids for the expression of guide RNAs and Dr. M. Pipkin (Scripps Research Institute) for providing pLMPd-Amt and pLMPd-GFP plasmids for expression of shRNAs. We acknowledge technical support from the following research cores at NYU Langone Health: the Genome Technology Center (RRID: SCR_017929), the Metabolomics core (RRID: SCR_017935), the Cytometry and Cell Sorting Laboratory (RRID: SCR_017926), Experimental Pathology Research Laboratory (RRID: SCR_017928).

## Funding

This study was funded by NIH grants AI125997, AI137004, AI180128 (S.F.), GM79465 (C.J.C) and a Colton Center for Autoimmunity pilot grant (S.F.). Additional funding was provided by postdoctoral fellowships from the American Society of Hematology (L.N.) and the National Multiple Sclerosis Society (L.W.).

## Author contributions

L.N., L.W., M.J., and A.T.P. conducted experiments. L.N., L.W., M.J., M.M., A.T.P., R.R., and M.S. analyzed data or interpreted results. L.N., L.W. and S.F. designed experiments. L.N. and S.F. wrote the manuscript. All authors read and approved the final version of the manuscript.

## Competing interests

S.F. is a scientific cofounder and consultant of CalciMedica. The other authors declare no competing interests.

## Data and materials availability

All unique and stable reagents generated in this study are available from the corresponding author with a completed materials transfer agreement.

## SUPPLEMENTAL FIGURE LEGENDS

**Fig S1: Characterization of mice with T cell specific deletion of Slc31a1. A**: Representative flow cytometry histograms and quantification of CTR1 expression in T-WT versus T-CTR1-KO pT_H_17 cells at day 3 (n = 8, 3 independent experiments). **B-G**: characterization of naïve T-WT and T-CTR1-KO mice. **B**: Total cell number in thymus (n = 6, 2 independent experiments). **C**: Representative flow cytometry plots and quantification of double negative (DN; CD4^-^CD8^-^), double positive (DP; CD4^+^CD8^+^), single positive CD4 (SP4; CD4^+^CD8^-^), single positive CD8 (SP8; CD4^-^CD8^+^) cells in thymus (n = 9, 3 independent experiments). **D**: Representative flow cytometry plots and total numbers of CD4^+^ and CD8^+^ cell numbers and frequencies in lymph nodes (inguinal, axillary, brachial) and spleen (n = 9, 3 independent experiments). **E**: Representative flow cytometry plots and total numbers of naïve and effector CD4^+^ T cells in lymph nodes (LNs) and spleen (n = 9, 3 independent experiments). **F**: Representative flow cytometry plots and total numbers of CD25^+^FOXP3^+^ T_reg_ cells in thymus (n = 6, 2 independent experiments). **G**: Representative flow cytometry plots and total numbers of CD25^+^FOXP3^+^ T_reg_ cells in LNs and spleen (n = 6, 2 independent experiments). Statistical significance was calculated by unpaired Student’s *t* test (A-B, D, F-G) or two-way ANOVA (C, E).

**Fig S2: CTR1 regulates pT_H_17 polarization in vitro. A**: Representative flow cytometry histograms and quantification of surface CD25, CD44, CD69 in CD4^+^ T cells activated in vitro for 24 and 48 hours (n = 6, 2 independent experiments). **B**: Representative flow cytometry histograms for RORγt in T-WT and T-CTR1-KO homT_H_17 cells, T-WT iT_reg_ cells shown as negative control, quantified in Figure 2C. **C**: Representative flow cytometry histograms for T-bet in T-WT and T-CTR1-KO T_H_1 cells, T-WT T_H_2 cells shown as negative control (left) and GATA3 in T-WT and T-CTR1-KO T_H_2 cells, T-WT T_H_1 cells shown as negative control (right); quantified in Figure 2D. **D**: Representative flow cytometry histograms for FOXP3 T-WT and T-CTR1-KO in iT_reg_, T-WT homT_H_17 shown as negative control; quantified in Figure 2E. **E**: Representative flow cytometry plots for GM-CSF production in T-WT and T-CTR1-KO pT_H_17 cells; quantified in Figure 2G. **F**: Representative flow cytometry plots for IL-17A and GM-CSF production in T-WT and T-CTR1-KO homT_H_17 cells; quantified in Figure 2H. **G**: Representative flow cytometry plots for IFN-γ production in T-WT and T-CTR1-KO T_H_1 cells; quantified in Figure 2I. **H**: Representative flow cytometry plots for IL-4 in T-WT and T-CTR1-KO T_H_2 cells, T-WT iT_reg_ shown as negative control; quantified in Figure 2I. Statistical significance was calculated by two-way ANOVA (A).

**Fig S3: CTR1 regulates pT_H_17 transcriptional program. A-F**: Bulk-RNAseq analysis of T-CTR1-KO versus T-WT CD4^+^ T cells cultured in either pT_H_17 or T_H_1 conditions in vitro for 3 days (n = 3). **A**: PCA comparing all conditions. **B**: Volcano plots of differentially expressed genes (DEGs) in T-CTR1-KO versus T-WT pT_H_17 (left) and T_H_1 (right) cells. **C**: Overlap of DEGs between pT_H_17 and T_H_1 culture conditions. **D**: IPA of DEGs for cells in pT_H_17 (left) and T_H_1 (right) cells. **E**: IPA upstream regulator analysis for DEGs in pT_H_17 (left) and T_H_1 (right) cells. **F**: Network clustering analysis for all DEGs in pT_H_17 cells. **G**: mRNA levels of pT_H_17 signature genes in T-WT pT_H_17 cells treated with 1 µM tetrathiomolybdate (TTM), 1 µM LCC-12 or control for 3 days, measured by RT-qPCR (n = 7, 2 independent experiments). **H**: mRNA levels of pT_H_17 signature genes in T-WT pT_H_17 cells treated with vehicle control, T-CTR1-KO pT_H_17 cells treated with 1 nM Elesclomol or vehicle control for 3 days, measured by RT-qPCR (n = 7-10, 2-3 independent experiments). Statistical significance was calculated by one-way ANOVA (G-H).

**Fig S4: CTR1 and copper regulate T cell metabolism. A**: ROS production in mitochondria measured by MitoSOX normalized over MitoView, in T-WT and T-CTR1-KO T_H_1 cells at day 3 (n = 3). **B**: Quantification of baseline OCR in T-WT and T-CTR1-KO T_H_1 cells at day 3 measured by Seahorse (n = 3). **C**: Quantification of basal and max OCR in T-WT and T-CTR1-KO pT_H_17 cells at day 1 (n = 6, 2 independent experiments). **D**: Quantification of basal and max OCR in T-WT and T-CTR1-KO T_H_1 cells at day 1 (n = 6, 2 independent experiments). **E**: Volcano plot of differential abundance of metabolites in T-CTR1-KO versus T-WT pT_H_17 cells at day 3, measured by LC-MS (n = 3). **F**: Glucose-dependence of T-WT and T-CTR1-KO pT_H_17 cells at day 3, measured by SCENITH assay (n = 5, 2 independent experiments). **G**: ATP production rate (pmol/min) in T-WT and T-CTR1-KO pT_H_17 cells at day 3 measured by Seahorse (2 independent experiments). **H**: *Sord* mRNA expression levels measured by bulk RNAseq in T-WT and T-CTR1-KO pT_H_17 and T_H_1 cells at day 3. **I**: Time course and quantification of OCR in T-WT pT_H_17 cells treated with 0.4 nM oligomycin or vehicle control, and T-CTR1-KO pT_H_17 cells treated with vehicle control for 3 days, measured by Seahorse. Basal respiration is quantified as a mean of 0-20 min, Max respiration is quantified as a mean of 50-70min (n = 3). **J**: Representative flow cytometry plots for T-bet (left) and IFN-γ (right); quantified in Figure 4I. Statistical significance was calculated by unpaired Student’s *t* test (A-F) or one-way ANOVA (G).

**Fig S5: CTR1 does not regulate pT_H_17 function via SOD1, ULK1 or MEK1/2. A**: SOD enzyme activity in CD4^+^ T cells activated for 3 days (n = 6, 2 independent experiments). **B**: Representative flow cytometry histograms and quantification of total ROS production in T-WT and T-CTR1-KO pT_H_17 and T_H_1 cells measured by DCFDA (n = 3-4, representative experiment). **C-D**: T-WT pT_H_17 cells treated with 150 nM LCS-1 or vehicle control for 3 days. Representative flow cytometry histogram and quantification of RORγt expression (**C**) (n = 9, 3 independent experiments). Representative flow cytometry plots of IL-17A and quantification of IL-17A and GM-CSF production after PMA/ionomycin stimulation (**D**) (n = 9, 3 independent experiments). **E-F**: T-CTR1-KO pT_H_17 cells treated with 25 µM Tiron or vehicle control for 3 days. Representative flow cytometry histogram and quantification of RORγt expression (**E**) (n = 6, 2 independent experiments). Representative flow cytometry plots of IL-17A and quantification of IL-17A and GM-CSF production after PMA/ionomycin stimulation (**F**) (n = 6, 2 independent experiments). **G**: Representative immunoblots and quantification of LC3 II normalized on βactin in T-WT and T-CTR1-KO pT_H_17 cells at day 3 (n = 3). **H-I**: T-WT pT_H_17 cells treated with 1 µM chloroquine (CQ) or vehicle control for 3 days. Representative flow cytometry histogram and quantification of RORγt expression (**H**) (n = 9, 3 independent experiments). Representative flow cytometry plots of IL-17A and quantification of IL-17A and GM-CSF production after PMA/ionomycin stimulation (**I**) (n = 9, 3 independent experiments). **J**: Representative immunoblots and quantification of p-ERK (Thr202/Tyr204) normalized on βactin in T-WT and T-CTR1-KO naïve cells with or without 10 minutes of aCD3 stimulation (n = 6-7, 2 independent experiments). Statistical significance was calculated by unpaired Student’s *t* test (A, G), paired Student’s *t* test (C-F, H-I) or two-way ANOVA (B, J).

**Fig S6: DNA methylation and histone acetylation regulate pT_H_17 polarization. A-B**: T-WT pT_H_17 cells treated with 200 µM fumarate, 100 µM succinate or vehicle control for 3 days. Representative flow cytometry histograms and quantification of RORγt expression (**A**) (n = 7, 2 independent experiments). Representative flow cytometry plots and quantification of IL-17A production after PMA/ionomycin stimulation (**B**) (n = 7, 2 independent experiments). **C**: Representative flow cytometry plots for T-bet and IFN-γ in T-WT T_H_1 cells treated with 200 µM 2-HG or vehicle control for 3 days; quantified in Figure 5D. **D**: Correlation analyses between differential 5mC and mRNA expression levels for key pT_H_17 signature genes shown in Figure 5G. **E**: Correlation analysis between differential 5mC and chromatin accessibility (ATACseq) for enhancer regions. **F**: mRNA levels of pT_H_17 signature genes in T-WT pT_H_17 cells treated with 25 µM TETi76 or vehicle control for 3 days, measured by RT-qPCR (n = 7, 2 independent experiments). **G**: Representative flow cytometry plots for T-bet and IFN-γ in T-WT T_H_1 cells treated with 25 µM TETi76 or vehicle control for 3 days; quantified in Figure 5J. **H**: mRNA levels of pT_H_17 signature genes in T-CTR1-KO pT_H_17 cells treated with 1 µM 5-Azacytidine or vehicle control for 3 days (n = 7, 2 independent experiments). **I**: RORγt expression shown as relative MFI normalized to control per experiment, and IL-17A production in T-WT pT_H_17 cells treated with 10 µM anacardic acid, 4 µM CPTH2, 100 nM A485 or vehicle control for 3 days. **J**: RORγt expression shown as relative MFI normalized to control per experiment, and IL-17A production in T-CTR1-KO pT_H_17 cells treated with 20 nM Trichostatin A (TSA) or vehicle control for 3 days, measured by flow cytometry (n=8, 3 independent experiments). **K**: RORγt expression shown as relative MFI normalized to control per experiment, and IL-17A production in T-CTR1-KO pT_H_17 cells treated with 50 nM SAHA (Vorinostat) or vehicle control for 3 days, measured by flow cytometry (n=10, 3 independent experiments). Statistical significance was calculated by unpaired Student’s *t* test (A-B, J, K), one-way ANOVA (C-D, I) or paired Student’s *t* test (F-H, J-K).

**Fig S7: deletion of CTR1 leads to a depletion of AP-1 motif binding in pT_H_17 cells. A**: Top motifs, TOBIAS analysis of ATACseq data (Figure 6B)**. B**: Top motifs, HOMER analysis of ATACseq data (Figure 6D).

**Fig S8: CTR1 is essential for pathogenicity of pT_H_17 cells in EAE. A**: *Slc31a2*, *Atp7a*, *Cd44* mRNA expression in donor 2D2 pT_H_17 cells isolated from spleen and spinal cords of mice with EAE, measured by bulk RNAseq (n = 3). **B**: *Slc31a2*, *Atp7a*, *Cd44* mRNA expression in blood mTeff from healthy control (HC) and patients with active relapsing-remitting multiple sclerosis (RRMS) measured by bulk RNAseq (GSE209596). **C-F**: T cells isolated from spinal cords of T-WT and T-CTR1-KO mice with active EAE (n = 9, 2 independent experiments). **C**: Representative flow cytometry plots and quantification of GM-CSF production after PMA/ionomycin stimulation. **D**: IL-2 production after PMA/ionomycin stimulation. **E**: Representative flow cytometry plots and quantification of CD25^+^FOXP3^+^ T_reg_ cells. **F**: IL-10 production after PMA/ionomycin stimulation. **G-H**: Passive EAE with shControl or sh*Slc31a1* 2D2 pT_H_17 cells injected in *Rag1* KO host mice. **G**: EAE disease score and incidence (n = 7, 2 independent experiments). **H**: Number of immune cells in spinal cords and spleen (n = 7, 2 independent experiments). **I-J**: Passive EAE with sgControl or sg*Slc31a1* Cas9^GFP^ 2D2 pT_H_17 cells injected in *Rag1* KO host mice. **G**: EAE disease score and incidence (n = 10, 3 independent experiments). **H**: Number of CD4^+^ T cells in spinal cords and spleen (n = 10-9, 3 independent experiments). Statistical significance was calculated by Welch’s *t* test (A, B), Mann-Whitney test (B), unpaired Student’s *t* test (C-F, H, J) or two-way ANOVA (G, I).

**Fig S9: Titrations for chemicals used in this study** Relative cell viability (top) and cell size (FSC-A MFI) measured in CD4^+^ T cells 3 days after activation and expansion in the presence of increasing concentrations of compounds and vehicle control. Grey box shows limiting concentration. Concentrations retained for the study are shown in green. **A**: TTM. **B**: LCC12. **C**: CuGTSM. **D**: Elesclomol. **F**: LCS-1. **G**: Tiron. **H**: CQ. **I**: 2-HG. **J**: succinate.

**Fig S10: Titrations for chemicals used in this study** Relative cell viability (top) and cell size (FSC-A MFI) measured in CD4^+^ T cells 3 days after activation and expansion in the presence of increasing concentrations of compounds and vehicle control. Grey box shows limiting concentration. Concentrations retained for the study are shown in green. **A**: fumarate. **B**: TETi76. **C**: Decitabine. **D**: 5-azacytidine. **F**: SAHA. **G**: Anacardic A. **H**: CPTH-2. **I**: A-485.

