## Supplementary figures and images for "CTR1-mediated copper uptake orchestrates metabolic-epigenetic regulation of pathogenic T_H_17 cells in autoimmune disease"

### Supplemental Figures

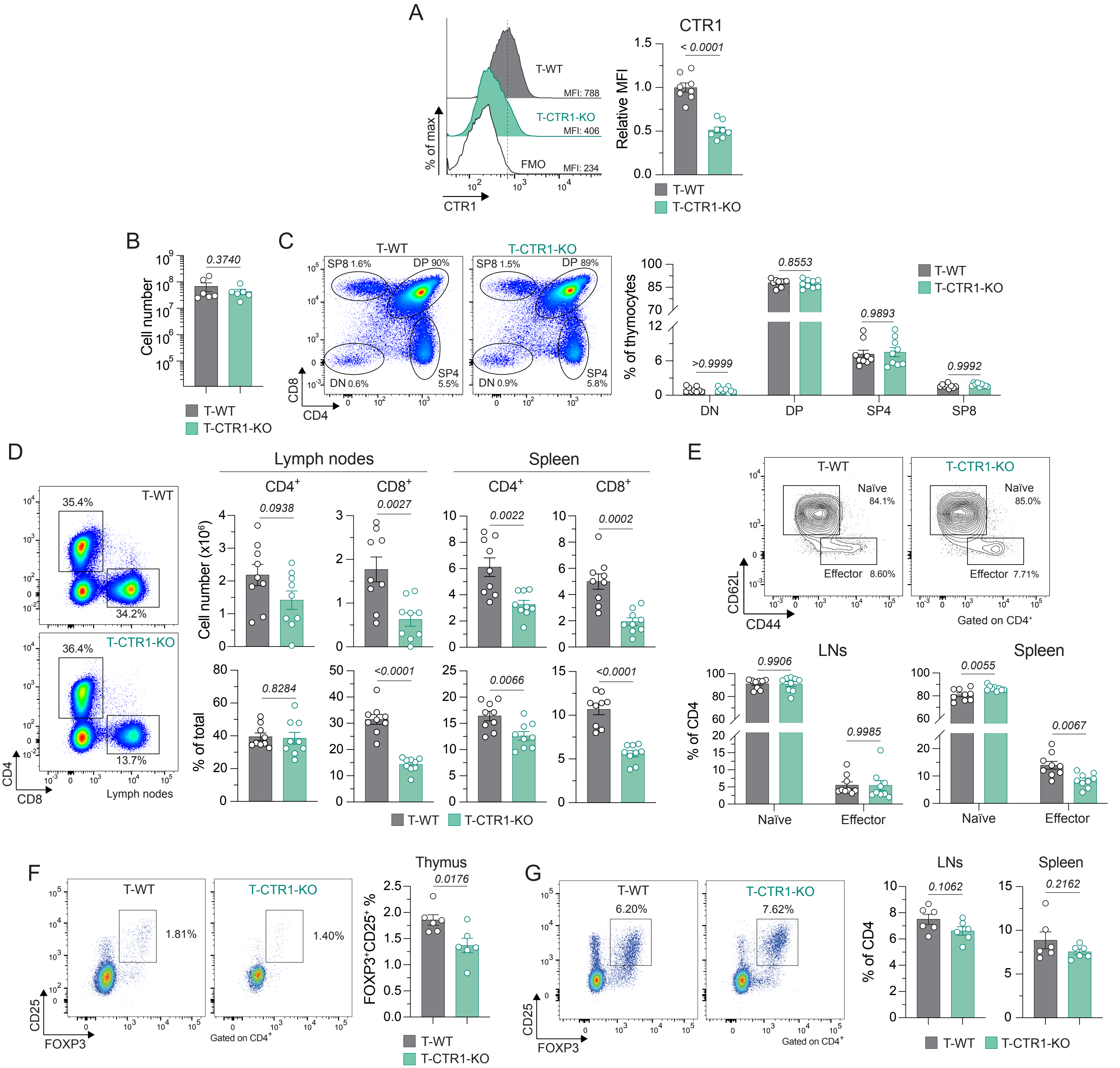

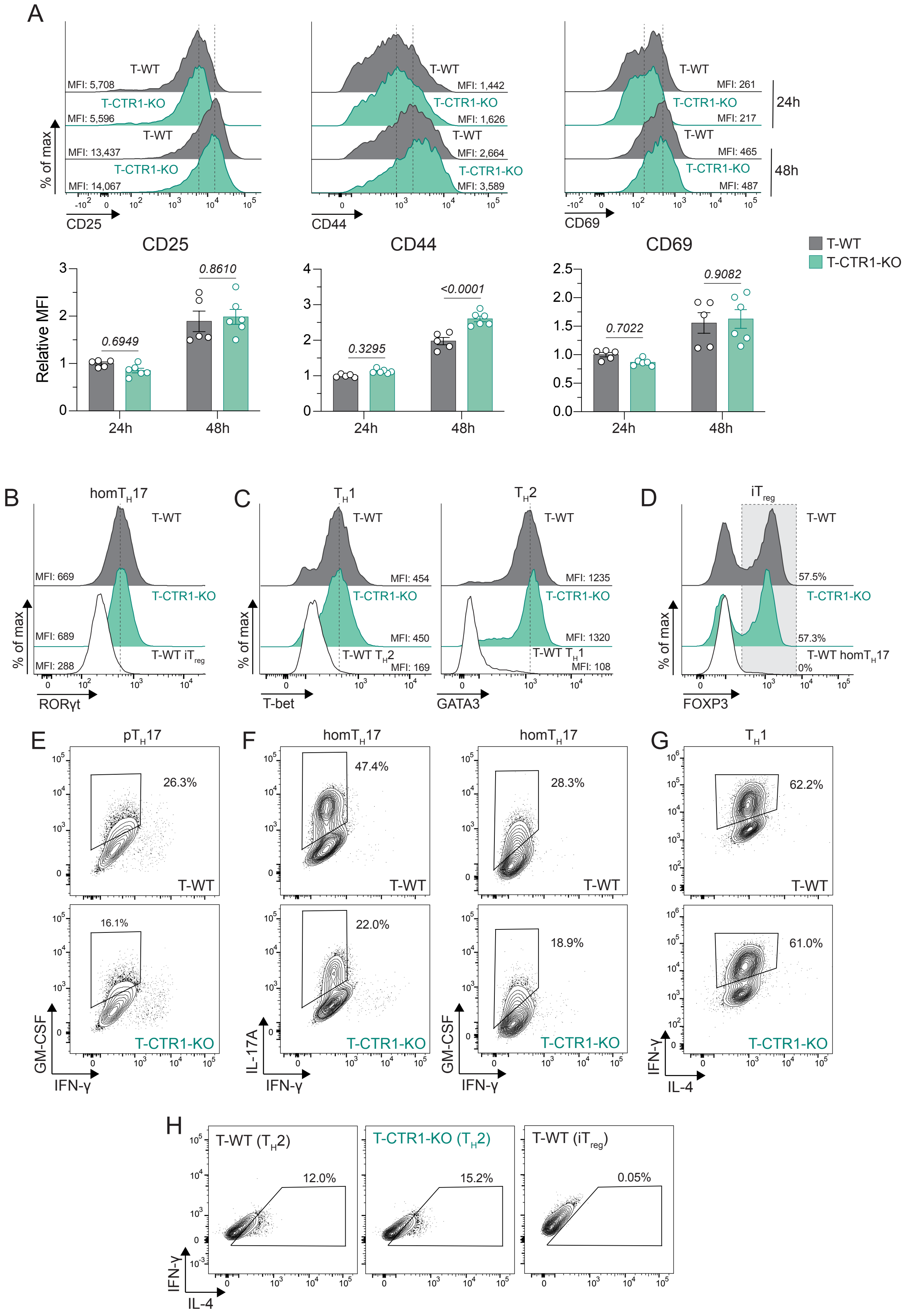

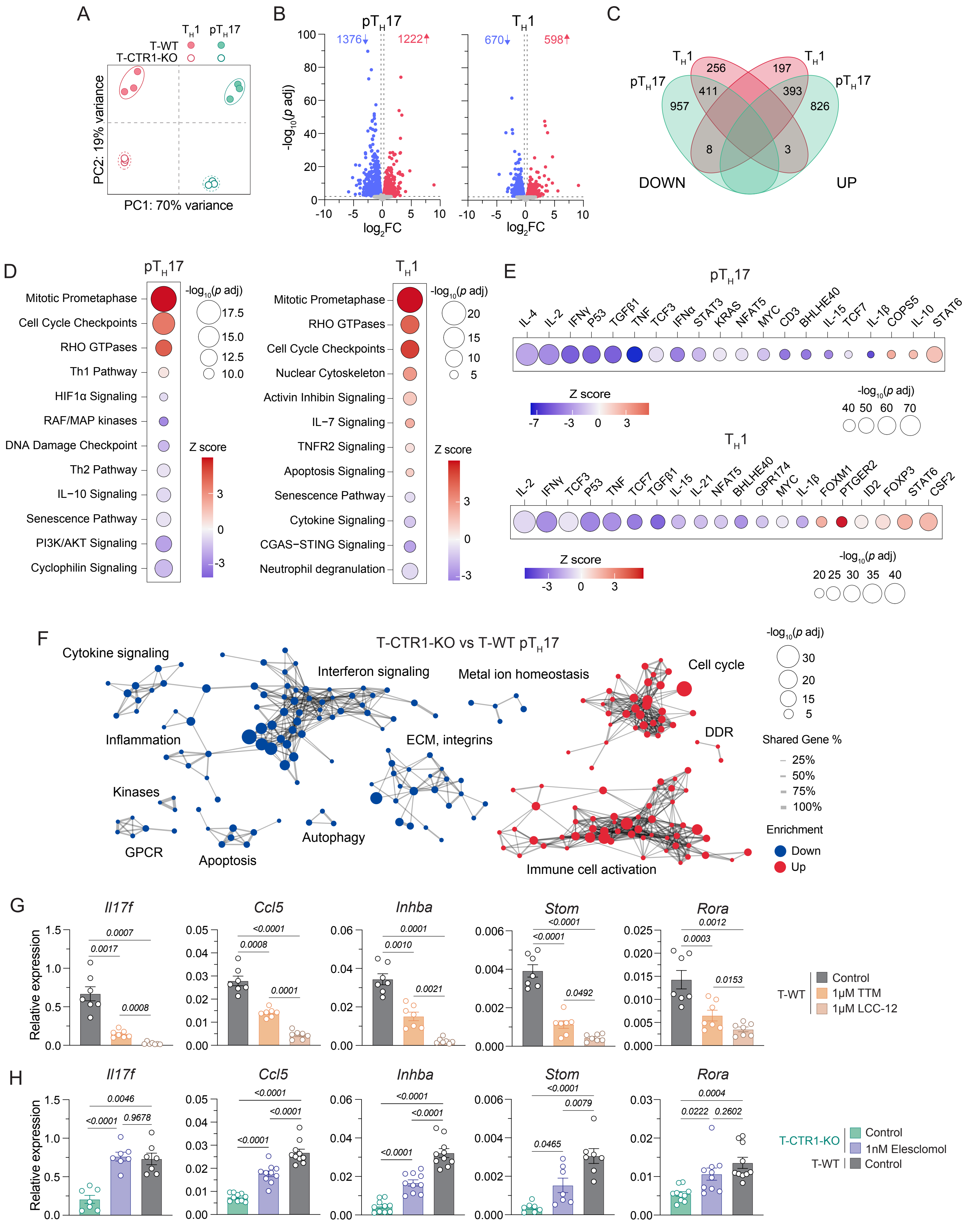

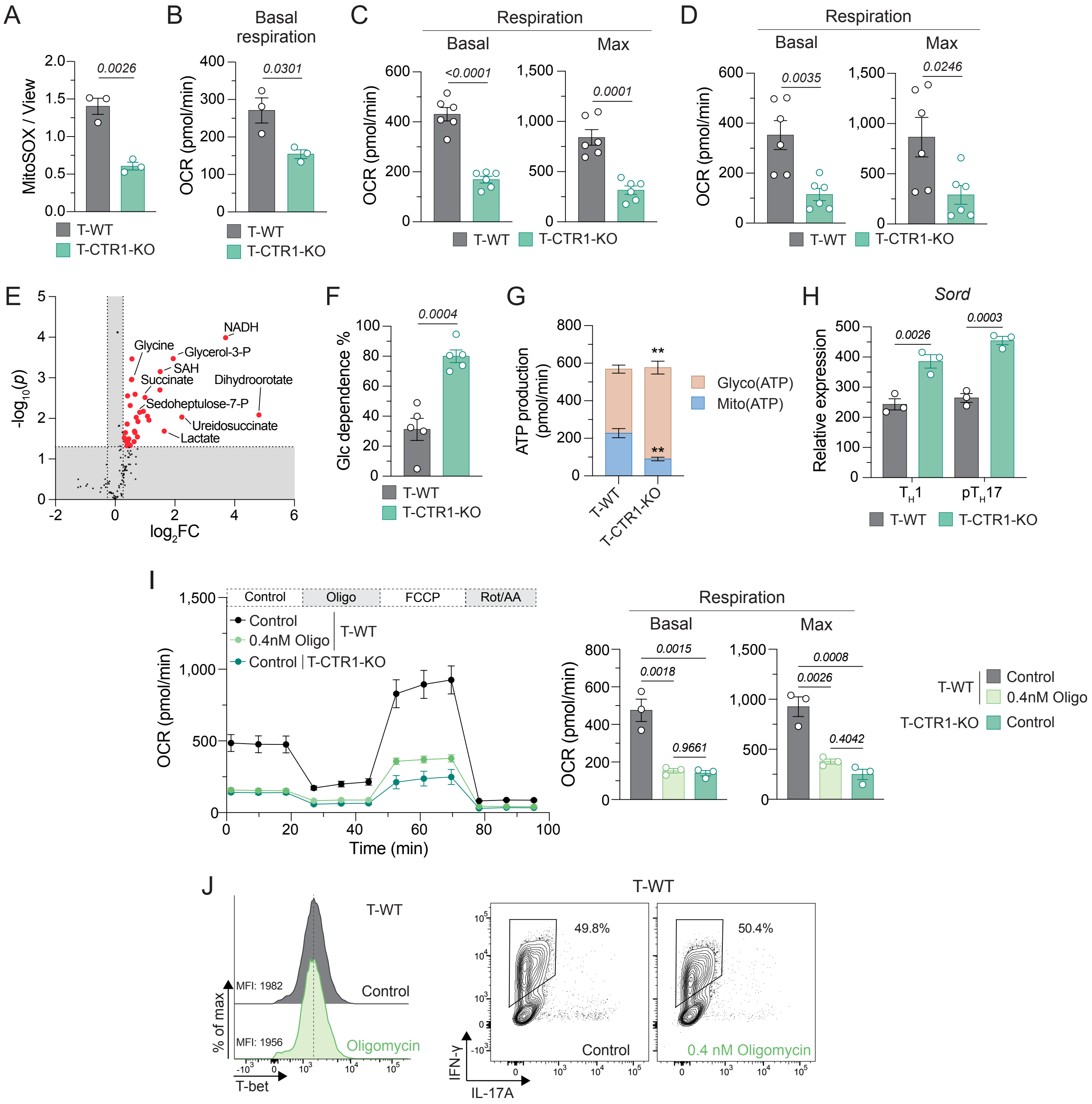

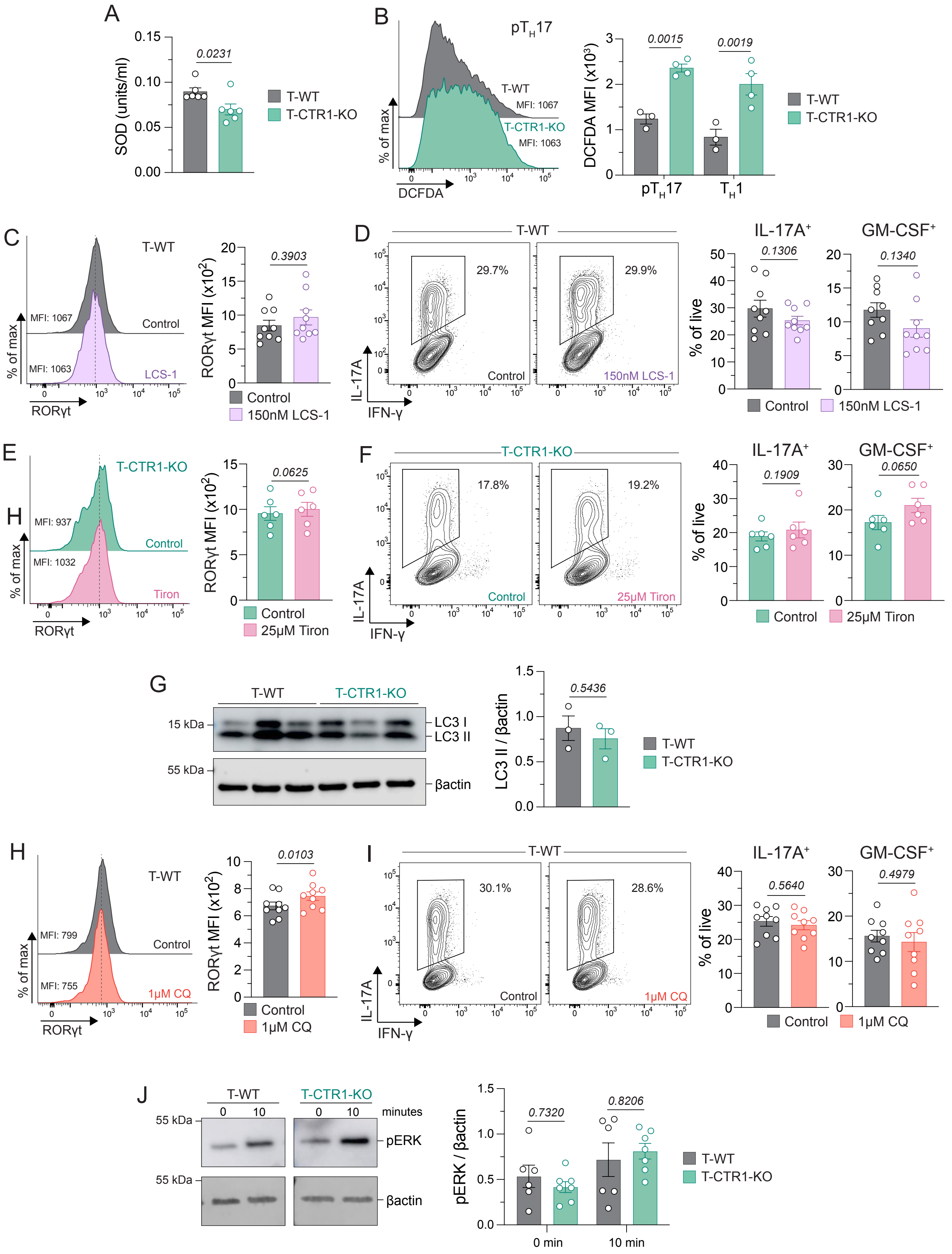

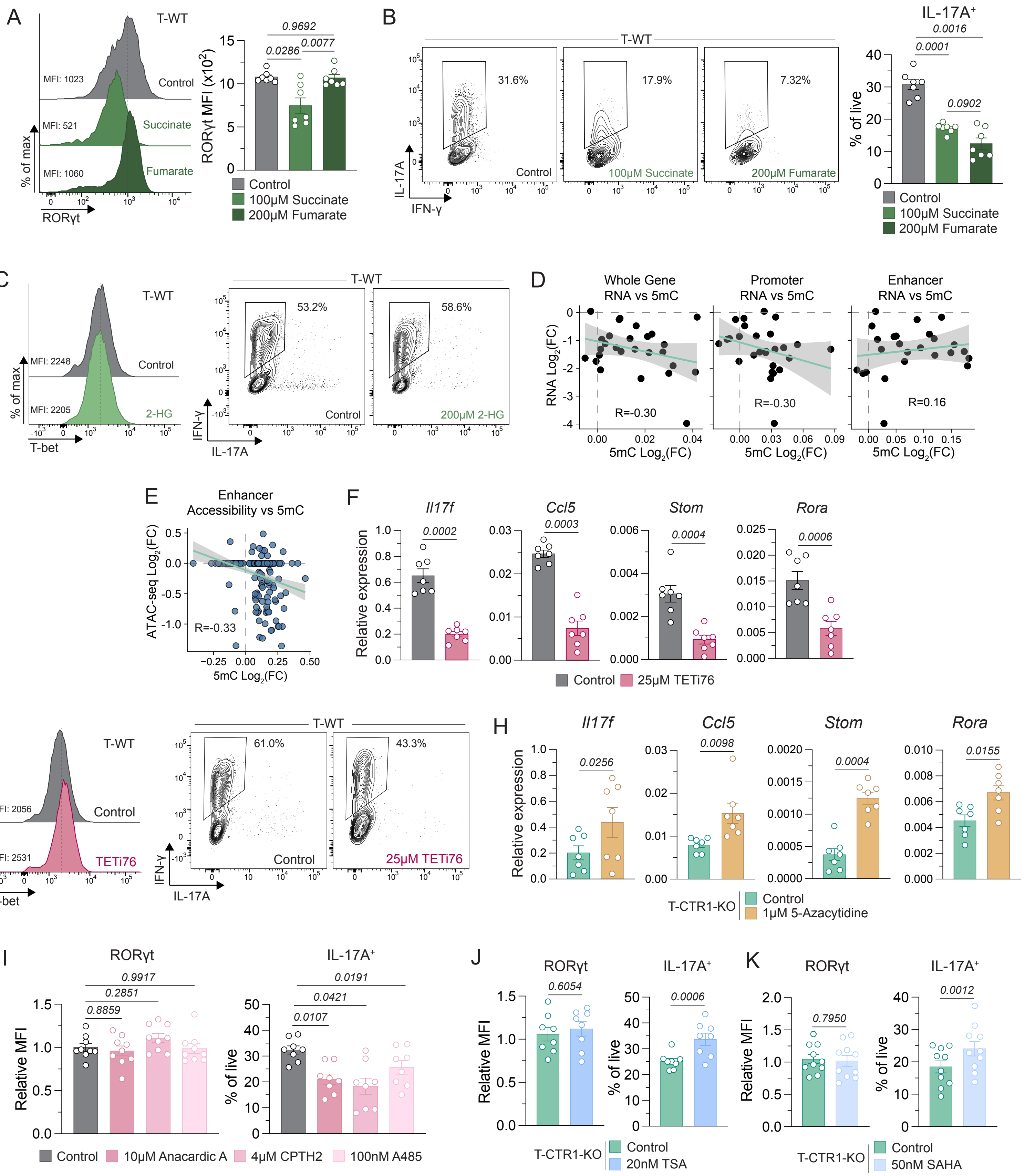

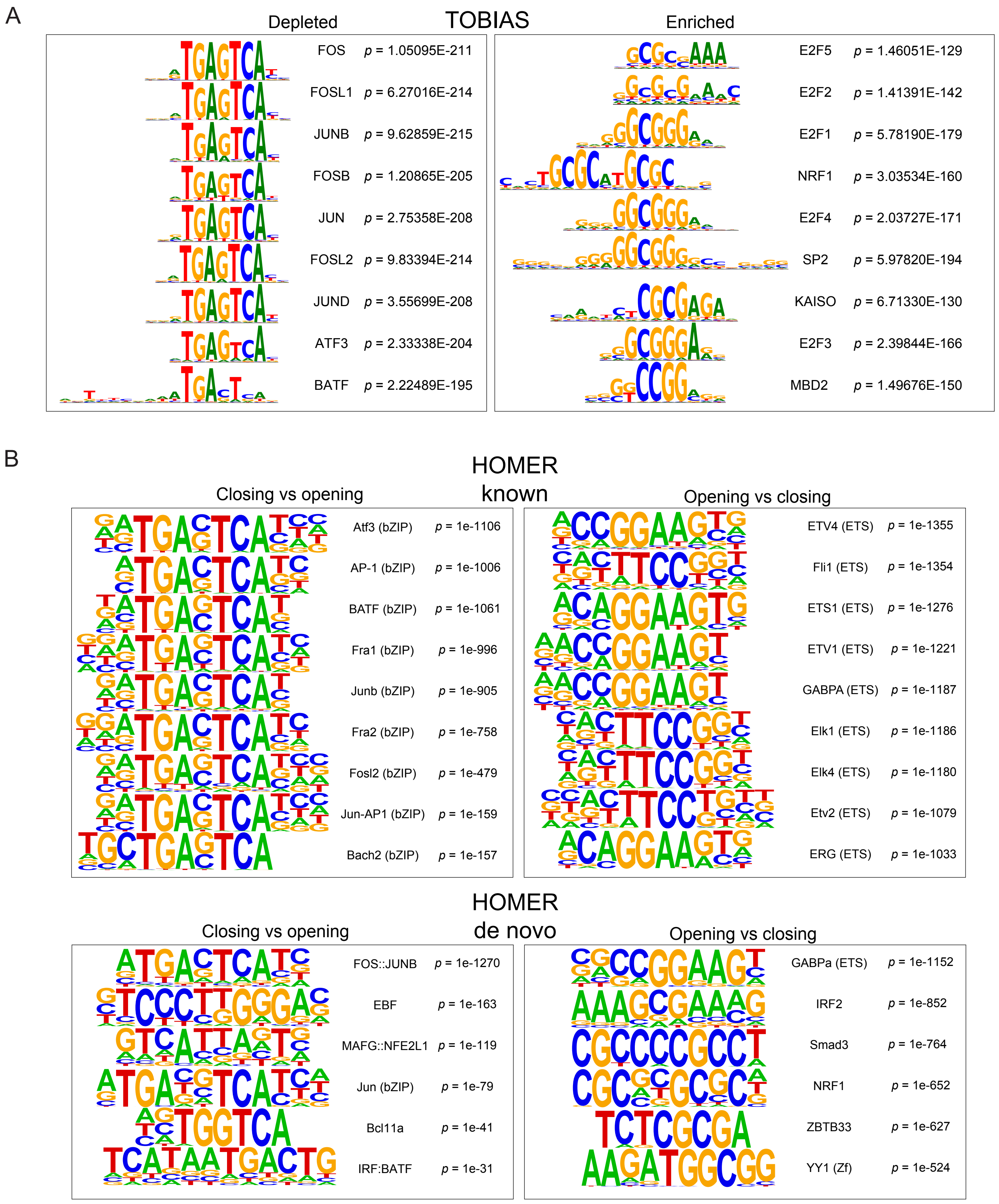

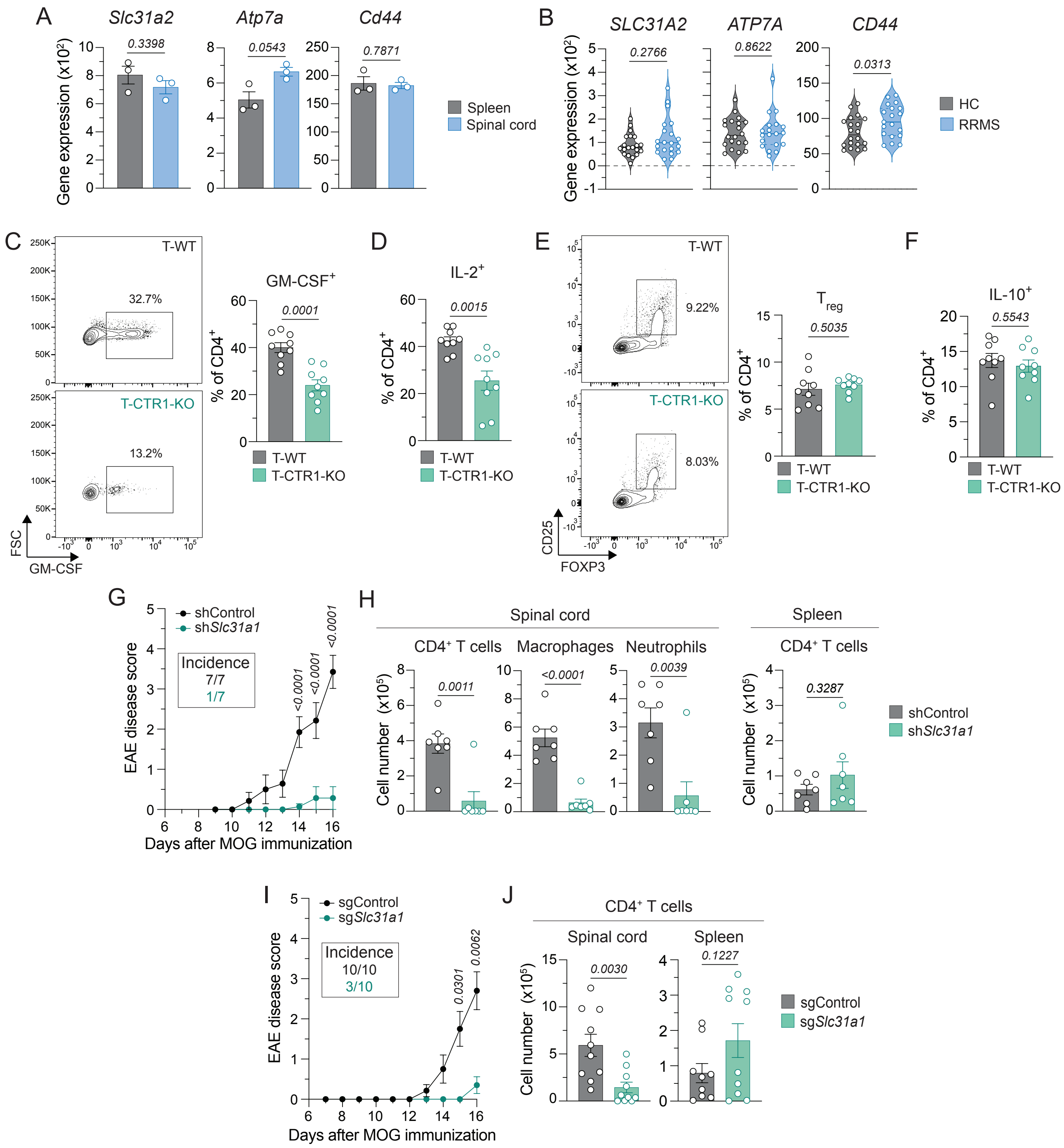

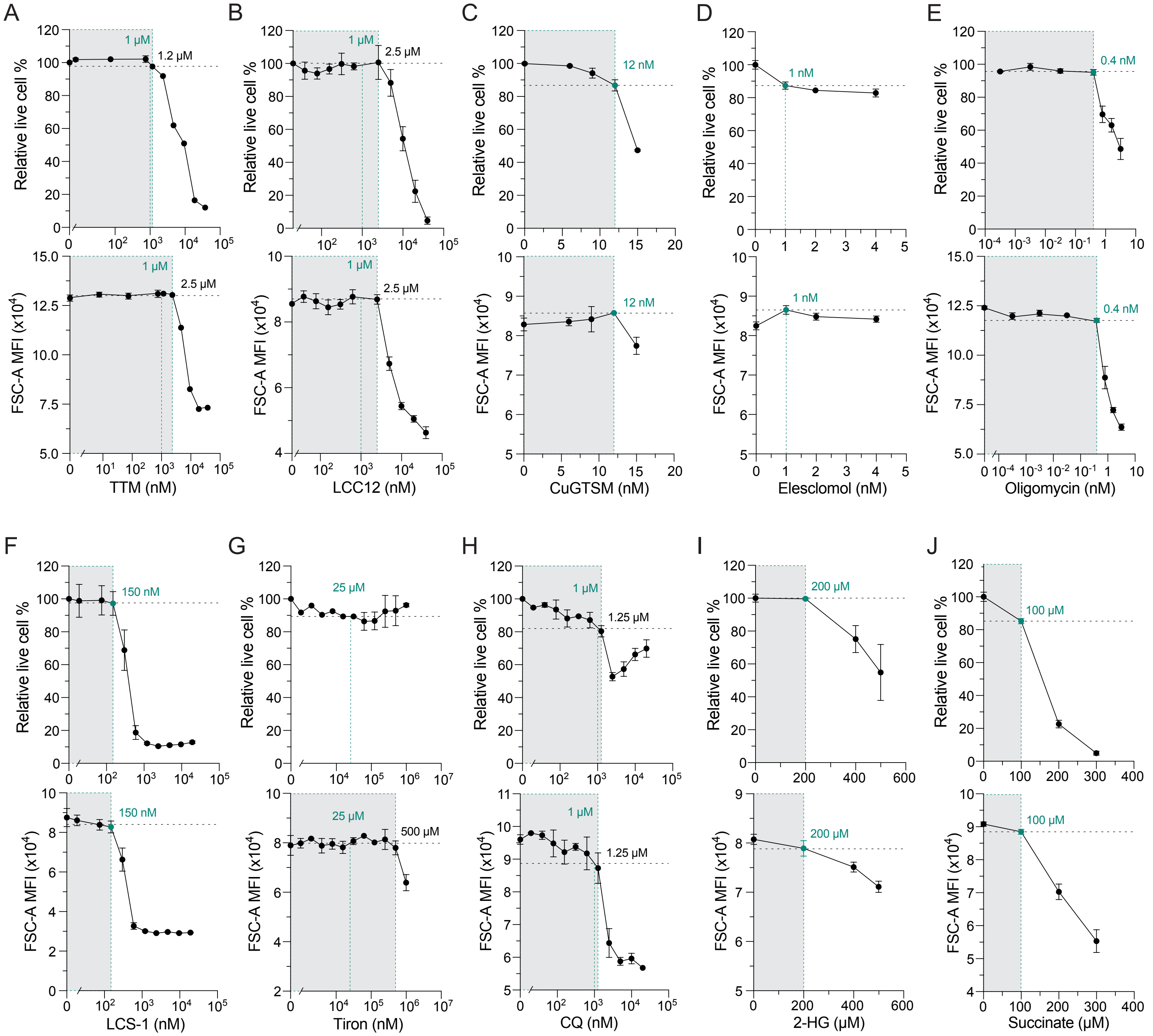

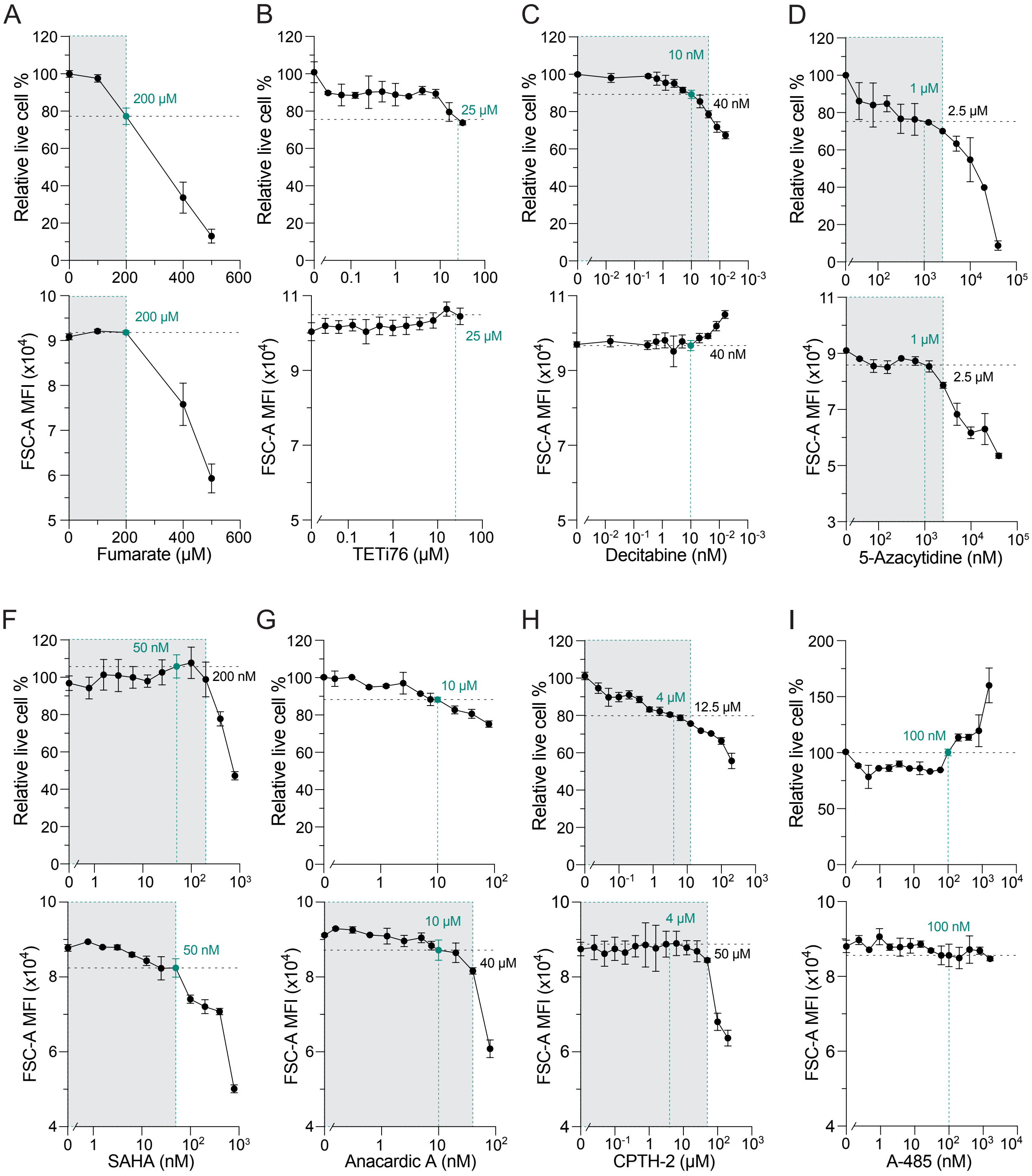
